# Minutes-timescale 3D isotropic imaging of entire organs at subcellular resolution by content-aware compressed-sensing light-sheet microscopy

**DOI:** 10.1101/825901

**Authors:** Chunyu Fang, Tingting Chu, Tingting Yu, Yujie Huang, Yusha Li, Peng Wan, Wenyang Feng, Xuechun Wang, Wei Mei, Dan Zhu, Peng Fei

## Abstract

Instant 3D imaging of entire organs and organisms at cellular resolution is a recurring challenge in life science. Here we report on a computational light-sheet microscopy able to achieve minute-timescale mapping of entire macro-scale organs at high spatial resolution, thereby overcoming the throughput limit of current 3D microscopy implementations. Through combining a dual-side confocally-scanned Bessel light-sheet illumination which provides thinner-and-wider optical sectioning of deep tissues, with a content-aware compressed sensing (CACS) computation pipeline which further improves the contrast and resolution based on a single acquisition, our method yields 3D images with high, isotropic spatial resolution and rapid acquisition improved by two-orders of magnitude. We demonstrate the imaging of whole brain (∼400 mm^3^), entire gastrocnemius and tibialis muscles (∼200 mm^3^) of mouse at subcellular resolution (0.5-μm isovoxel) and ultra-high throughput of 5∼10 minutes per sample. Various system-level cellular analyses, such as mapping cell populations at different brain sub-regions, tracing long-distance projection neurons over the entire brain, and calculating neuromuscular junction occupancy across whole muscle, were also readily enabled by our method.

## Introduction

The comprehensive understanding of majority and singular cellular events, as well as their complex connections in whole organs and organisms, is one of the fundamental quests in biology. To extract the various cellular profiles of different physiological functions within specimens, three-dimensional (3D) high-resolution imaging is required throughout a mesoscale-sized volume. However, creating such a large-scale image dataset has posed significant challenges for current 3D light microscopy methods, which show relatively small optical throughputs^1,2^. Furthermore, in thick mammalian organs, severe light scattering and attenuation greatly limit the complete extraction of signals from deep tissues. A common strategy for addressing these issues has been to use 3D tile stitching combined with tissue sectioning^2-5^. For example, both tiling confocal microscopy^6-9^ and sequential two-photon tomography^10,11^ (STPT) can three-dimensionally image mouse brain at subcellular resolution, but do so at the expense of a very long acquisition time due to the slow laser-point-scanning. The recent integration of light-sheet microscopy^12-15^ and tissue-clearing^16-19^ has facilitated an important alternative to conventional 3D histological imaging approaches by instead applying a nondestructive light-sheet to selectively illuminate a thin plane of the clarified sample. At the same time, the use of wide-field detection results in improved imaging speed as compared to the epifluorescence approaches. Although light-sheet microscopy using a Gaussian-type hyperbolic light-sheet^2,13,20^ has been successfully used for imaging large organ samples, a trade-off exists between the minimum thickness of the light-sheet and the confocal range over which it remains reasonably uniform. As a reference, when illuminating a range of 1 mm in length, an optimized Gaussian light-sheet diverges to a full-width at half-maximum (FWHM) thickness of 11 μm at either end, and is hence incapable of resolving fine neuronal fibers across a large volume of brain. Sweeping the light-sheet along the propagation direction can extend the confocal range, but the achieved axial resolution remains inadequate, and tissue scattering effects also limit its propagation range^21,22^. Bessel-type light-sheet microscopy^23-25^ (LSM) can generate a thin and non-diverging light-sheet that can illuminate samples with a much larger field of view (FOV) while maintaining high axial resolution. Considering the necessity of minimizing excitation from the side lobes of the Bessel beam^26,27^, Bessel LSM usually uses a high numerical-aperture (NA) detection objective with a small depth of focus to reduce the fluorescence excitation by the side lobes. Thus, it is mainly optimized for the imaging of organelles in a single or a few cells. However, as with epifluorescence methods that suffer from a trade-off between accuracy and scale, Gaussian and Bessel LSM systems still have limited optical throughput, which is far from adequate for obtaining high spatial resolution across a very large FOV^28^. To image a whole mouse brain at subcellular resolution, a small step size and tile-stitching at high magnification are necessary to acquire trillions of sub-μm^3^ voxels over a cm^3^-sized volume^2,19^. Therefore, a long acquisition time from several hours to days, as well as increased photobleaching to samples, still prevents the widespread applications of LSM to the high-resolution mapping of entire mammalian organs/organisms.

Here we show that combining a large-FOV scanning Bessel light-sheet microscopy with a post-computation procedure, termed content-aware compressed sensing (CACS), can address the aforementioned imaging challenges. A dual-side confocally-scanned Bessel light-sheet with a millimeter-to-centimeter tunable range is developed to illuminate regions-of-interest (ROIs) from a few cells to entire mouse organs, providing uniform optical sectioning of deep tissues with 1-5 μm ultra-thin axial confinement. In addition to the optical design, our content-aware compressed sensing (CACS) computation can further improve the contrast and resolution of the acquired 3D image based on a single input of its own, allowing satisfying spatial resolution with data acquisition shorten by two-orders of magnitude. Our method thus overcomes the limitations of anisotropic resolution as well as inadequate throughput from current whole-organ imaging techniques. We apply this method to the high-throughput anatomical imaging of whole mouse brain to demonstrate its unique capabilities, such as the rapid screening of multiple brains in minutes, instant imaging of any region of interest in selected brain at subcellular resolution in seconds, and teravoxel high-resolution mapping of entire brain of ∼400 mm^3^ on a timescale of ∼10 minutes. Such a readily-accessible whole brain dataset allows us to perform various system-level cellular profiling, such as segmentation of brain regions, cell population counting, and tracing of neural projections. Furthermore, we also demonstrate dual-color imaging of subcellular neuromuscular junctions across an ∼200 mm^3^ volume of entire mouse muscle in ∼5 minutes, thus enabling efficient quantification of the neuromuscular junction occupancy at whole tissue scale.

## Results

### Zoomable, line-synchronized, CACS Bessel plane illumination microscopy

The optical layout and photographs of our self-built Bessel light-sheet microscope are shown in **Supplementary Fig. 1 and 2**, respectively. The simple mode of operation involves sweeping the Bessel beam in the ***y*** direction to create a continuous sheet at each ***z*** plane. However, in this mode, the cross-sectional profile of the excitation sheet contains broad tails because of the combined influence of the side lobes^24^ (**Supplementary Fig. 3**). Given the fact that the increase in depth-of-field is proportional to the square of the decrease in NA, the axial excitation from these side lobes is more likely to deteriorate the fluorescence detection in our low-to-middle magnification/NA setup (**Supplementary Fig. 4**). A confocal slit was therefore formed by tightly synchronizing the sweeping Bessel beam with the rolling active pixel lines of the camera^29,30^, to block the influence of residing fluorescence excited by the side lobes, and led to much less background (**Supplementary Figs. 3, 4, Video S1)**. As a result, the microscopy system could produce sharp optical sectioning at a very large scale, for example, generating a 4 × 4 mm illuminating light-sheet with a cross-sectional profile of ∼2.7 μm in thickness, as compared with an over 20-μm thick Gaussian sheet covering the same FOV (**Supplementary Fig. 3, Video S1**). When this wide-and-thin Bessel plane illumination combined with a matched magnification (3.2×) was applied to the 3D imaging of cleared mouse brain tissue with neurons labelled by green fluorescent protein (Thy1-GFP-M), the maximum intensity projection (MIP) in the ***y-z*** plane of the cortical area (**Fig. 1c**) showed axial resolution and contrast superior to those from 3.2× epi-illumination and 3.2× Gaussian plane illumination (**Fig. 1a, b**).

**Fig. 1.**
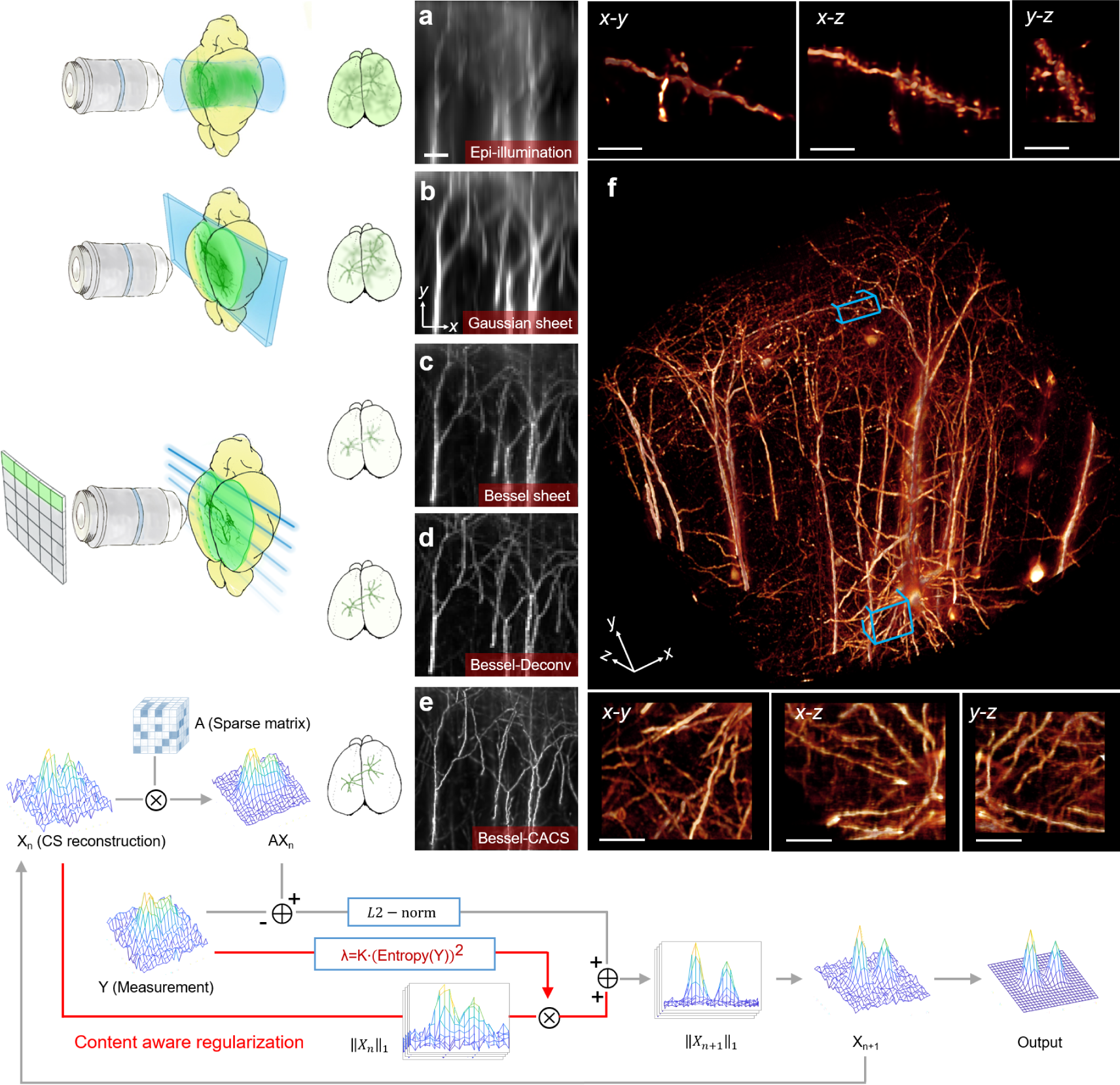
CACS Bessel plane illumination microscopy and other imaging modes. **a**, Epi-illumination mode (left), and maximum-intensity-projection (MIP) in the ***y-z*** plane from a 3D image stack of a cleared transgenic mouse brain tissue with GFP labelled to the neurons (right, 3.2×/0.28 objective). **b**, Gaussian sheet mode (left) showing plane illumination at selective depth of brain, and ***y-z*** plane MIP from the same sample (right, 0.02 NA illumination + 3.2×/0.28 detection). **c-e**, Raw, Deconvolution and CACS modes under our synchronized Bessel plane illumination geometry (0.14 NA illumination + 3.2×/0.28 detection), showing much sharper optical sectioning (**c, d**, left) and how CACS computation being further applied to super-resolve the image (**e**, left). The ***y-z*** plane MIPs by each mode are accordingly shown in right columns. Scale bar, 20 μm. **f**, Volume rendering of GFP-labelled neurons in a cleared mouse cortex tissue by 3.2× Bessel-CACS mode. Insets: MIPs along orthogonal axes of the cubical volume of interest (blue boxes). Scale bar, 10 μm.

The raw Bessel sheet working at such a relatively large FOV permits rapid 3D imaging of whole brain without many times of stitching, but may still show inadequate resolutions, for example, ∼4 μm isotropic resolution under magnifications factor of 3.2× plus 2-μm ***z*** step size. Normally, the raw Bessel sheet data can be deconvolved with the appropriate point spread functions (PSFs), to slightly improve image quality (**Fig. 1d**). An effective means to improve the contrast and resolution of a single measurement is compressed sensing (CS) computation, which is well-known for its ability to recover signals from incomplete measurements^31^. In its conventional light microscopy implementation, CS has recovered degraded signals from camera pixilation (wide-field) or incomplete scanning (confocal), and reduced the acquisition time for each 2D frame^32,33^. Combining with light-sheet microscopy, CS increased the SNR of images, thereby the acquisition time was decreased^34,35^. It was also used to improve signal quality for 2D speckle image in nonlinear structured illumination microscopy^36^. In our Bessel light-sheet setup, the camera’s under-sampling in the ***x-y*** plane and sparse ***z***-excitation by thin plane illumination allows the application of compressed sensing. To adopt this approach to Bessel imaging, in which the signal sparsity and dynamic range may vary drastically within a large volume^37^ (**Supplementary Figs. 6, 7**), we extended the CS computation to three dimensions and applied content-aware (CA) regularization that was highly adaptive to the signal characteristics (**Fig. 1e**). Using our CACS processing, we aimed to achieve a higher-quality image ***x*** from the raw large-scale Bessel image ***y***, which shows inadequate resolution resulting from the voxelization and limited numerical aperture. We first obtained the compressed sensing matrix ***A*** in sparse domain based on the system PSF, and applied it to the conventional CS computation term. For the content-aware regularization, the entire Bessel image stack ***y*** was first divided into multiple small volumes ***y***_***i***_ to calculate the specific entropy parameter ***β***_***i***_ that represents the degree of signal disorder in each ***y***_***i***_ (**Supplementary Note 3**). A content-aware regularization factor ***λ***_***i***_ is then determined from the calculated ***β***_***i***_, as well as a signal density index ***α***_***i***_, and applied to the weighting of regularization term ‖*x*_*i*_‖_1_, which is the ***L1***-norm of the ***x***_***i***_ to be solved. This content-aware regularization with appropriate ***λ***_***i***_ balances the iterative signal reconstruction process between an overfitting result subject to excessive constraints from complicated signals and an under-fitting result with too sparse signals accompanied by obvious artefacts that are difficult to further optimized by iteration (**Supplementary Note Fig. 1, Video S2**). A higher-resolution image tile ***x***_***i***_ can be recovered by finding the optimal solution of the following problem in the sparse Fourier domain:

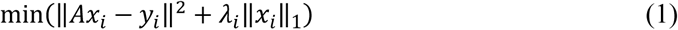

Finally, the multiple small CACS reconstruction ***x***_***i***_ were stitched together to form the final high-resolution large-scale output (**Supplementary Note Fig. 1, Video S2**). **Fig. 1e** shows the super-resolved ***y-z*** MIP of the neurons originally by 3.2× Bessel sheet, after a 4-times CACS computation applied. **Fig. 1f** shows a CACS-reconstructed volume rendering of apical dendrites in a region of the cortex yielding sufficiently high isotropic 3D resolution and signal-to-noise ratio (SNR) to discern subcellular structures such as dendrite spines. Besides the recovery of signals originally imaged at relatively low magnification, CACS procedure was also applicable to higher magnifications, such as 12.6×, for resolving ultra-fine dendrite spines at deep subcellular resolution (**Supplementary Fig. 8**).

### Comparisons with Gaussian sheet, confocal and Two-Photon-Excitation microscopy

As confocal microscopy remains the workhorse for 3D cellular imaging and the Gaussian light-sheet method is the standard for light-sheet microscopy, we compared both techniques with various modes of Bessel beam plane illumination. Image slices in the ***x-z*** plane from dendrites of Thy1-GFP-M mouse brain demonstrating comparative axial resolutions were acquired using a point-scanning confocal microscope (Nikon Ni-E, CFI LWD 16×/0.8W objective), a two-photon-excitation (TPE) microscope (Nikon Ni-E, CFI LWD 16×/0.8W objective), a selective plane illumination microscope (SPIM, 0.02 NA illumination, 3.2×/0.28 detection objective), the low-resolution Bessel sheet mode (0.14 NA illumination + 3.2×/0.28 detection), the high resolution Bessel sheet mode (0.28 NA illumination + 12.6×/0.5 detection objective) and the CACS-enabled Bessel sheet mode (**Fig. 2a-f**). With the confocal and TPE microscopes, the anisotropic PSFs in the epifluorescence mode influenced the visualization of the dendrite fibers in the longitudinal direction (**Fig. 2a, b**). The thick Gaussian sheet covering the full imaging FOV (∼4 mm) also gave insufficient axial resolution (FWHM ∼15 μm, **Fig. 2c**), which was obviously poorer than that of the other methods. The raw 3.2× Bessel sheet covering the same FOV gave higher axial resolution superior to the Gaussian sheet (**Fig. 2d**). In contrast, the 12.6× Bessel sheet mode and CACS reconstruction both demonstrated a clear reduction in out-of-focus haze, as well as high axial resolution superior to the Gaussian sheet, 3.2× Bessel sheet, confocal and TPE methods (**Fig. 2e, f**). The CACS reconstruction substantially recovered the ultrastructures of the dendrites (**Fig. 2f**), which remained unresolvable in the raw 3.2× Bessel sheet image (**Fig. 2d**). In the meantime, the recovery fidelity was shown to be sufficiently high, as compared to the 12.6× Bessel sheet image (**Fig. 2g, h**). The line cuts through individual dendrite fibers (**Fig. 2a-f**) made by each method further reveal the narrower axial and lateral FWHMs of the CACS Bessel sheet modes in comparison with the regular Bessel sheet. The experimentally measured axial FWHMs were around 15, 4.5, 1.5, 1.5, 4.2 and 4 μm for Gaussian sheet, 3.2× Bessel sheet, 12.6× Bessel sheet, 3.2× CACS Bessel sheet, confocal and TPE methods, respectively (**Fig. 2k**). The corresponding lateral values were around 4.5, 4.5, 1.5, 1.5, 0.85 and 0.85 μm. Aside from the resolution improvement, the CACS procedure also significantly improved the SNR, as shown in **Supplementary Fig. 9**.

**Fig. 2.**
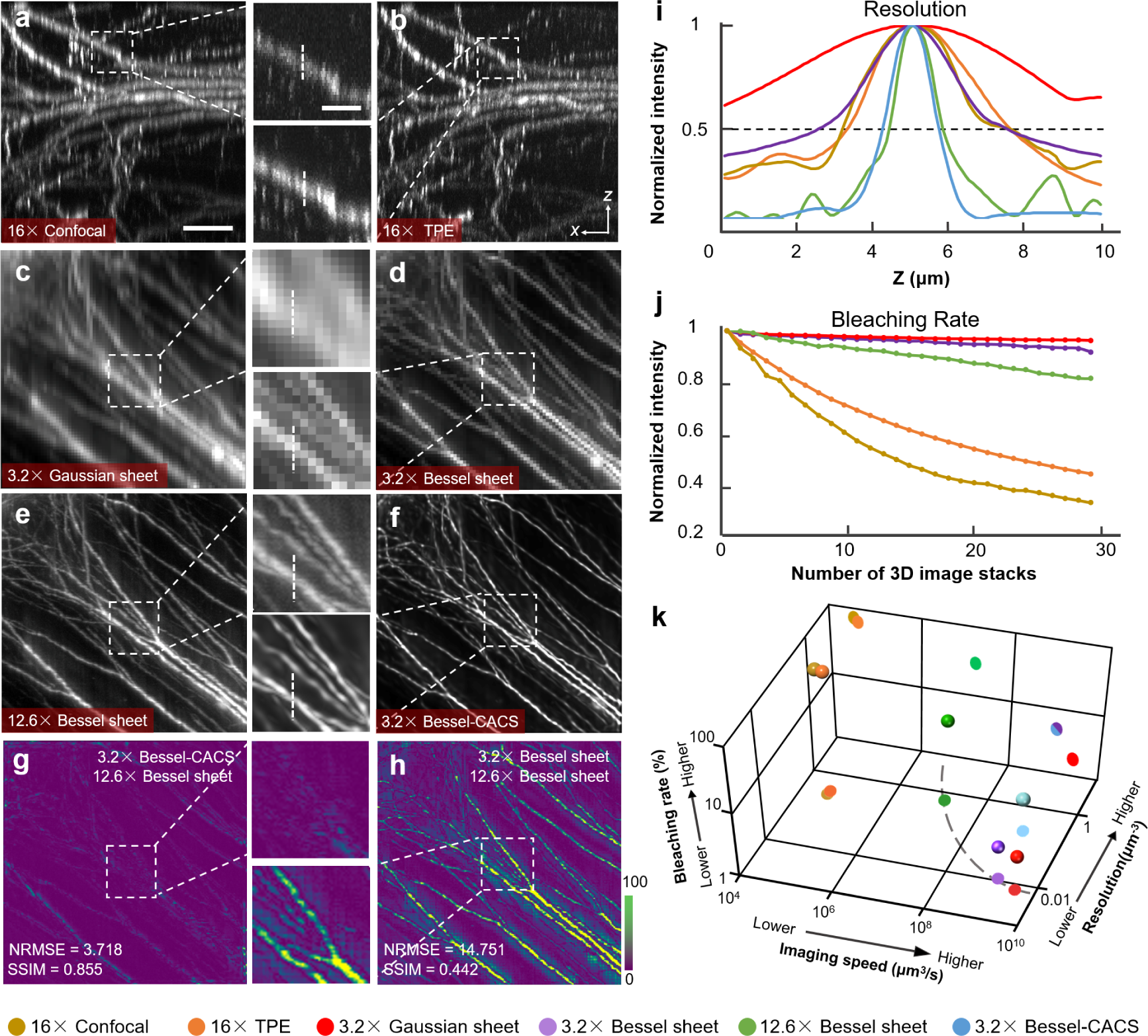
Comparisons of CACS Bessel sheet modes to confocal, TPE, and SPIM. **a-f**, Comparative image slices in an ***x-z*** plane through GFP-labelled neurons in a cleared mouse cortex by: point-scanning confocal microscopy (**a**, 16×/0.8); two-photon excitation microscopy (**b**, 16×/0.8); Gaussian sheet (**c**, 0.02 NA illumination + 3.2×/0.28 detection); low-resolution Bessel light-sheet illumination (**d**, 0.14 NA illumination + 3.2×/0.28 detection); high-resolution Bessel light-sheet illumination (**e**, 0.28 NA illumination + 12.6×/0.5 detection); CACS Bessel sheet which successfully resolves the single neurons with using the same low-resolution setup (**f**, 0.14 NA illumination + 3.2×/0.28 detection). Scale bar, 30 μm (inset, 10 μm). **g**,**h**, Error maps of 3.2× Bessel-CACS output (**f**) and raw 3.2× Bessel sheet result (**d**), as compared with 12.6× Bessel sheet result (**e**). The calculated normalized root mean square error (NRMSE) and structural similarity (SSIM) further quantitatively confirm the high recovery fidelity of CACS as well as the resolution improvement. **i**, Linecuts (as shown in insets in **a-f**) for each method, with 50% intensity level shown for estimation of the FWHM. **j**, Bleaching rates obtained from repeated 3D imaging (30 times) of neurons in mouse cortex, normalized to account for the differences in SNR. **k**, 3D plot comparing the acquisition speed, bleaching rate and volumetric resolution of 6 imaging modes indicated by different colors. The dashed line that connects the data points of 3.2× Bessel sheet, 12.6× Gaussian sheet and 12.6× Bessel sheet projected in speed-resolution plane represents the throughput limit in conventional light-sheet microscopes.

For large-scale imaging, which often encompasses data from hundreds of gigavoxels, the achievable resolution needs to be carefully balanced with the imaging time, because of the limited space bandwidth product (SPB) of the system^28^. The achievement of sufficiently precise measurements in a reasonably short acquisition time with a low photo-bleaching rate thereby poses a big challenge. For example, confocal and TPE microscopes can three-dimensionally image a sample at fairly high resolution (3 μm^3^), but do so at the cost of low speed due to the point scanning detection (4.2×10^4^ μm^3^/s), and noticeable phototoxicity caused by the epi-illumination mode. In contrast, Gaussian or Bessel sheet modes with in-plane excitation and camera-based detection can image the sample at video rate speed and with lower photo-bleaching. The CACS-enabled Bessel sheet mode further improves the volumetric resolution by about 30 times, while maintaining the same acquisition speed and bleaching rate, thereby resulting in a much higher throughput.

We compared the degree of photo bleaching of the five imaging methods by repeatedly imaging the cortex area of a Thy1-GFP-M mouse brain 30 times (**Supplementary Fig. 10**). After normalizing for differences in SNR (see **Methods**), we calculated the averaged signal intensity of each volume for each method, and plotted their variations in **Fig. 2j**. After 30 cycles of imaging of the same volume, approximately 95%, 90%, 80%, 40%, and 30% of the fluorophores were preserved by the Gaussian sheet, 3.2× Bessel sheet, 12.6× Bessel sheet, TPE, and confocal methods, respectively. In **Fig. 2k** and **Supplementary Note Table 1**, we further rate the overall imaging performance of confocal, TPE, Gaussian sheet, 3.2× Bessel sheet, 12.6× Bessel sheet and CACS Bessel sheet methods by comparing their volumetric resolutions, acquisition speeds, and photo-bleaching rates. Under the magnification of 3.2× used for whole-brain imaging, the CACS Bessel sheet mode yielded an isotropic resolution of ∼1.5 μm at a high acquisition rate of ∼1.5 mm^3^ s^-1^. As shown in the data projected to a resolution-speed plane, the CACS Bessel sheet mode breaks the SPB limitation defined by the dashed resolution-speed curve. Thus, it can be considered to be a tool facilitating the combination of large FOV with isotropic high-resolution, which is difficult to meet using previous methods.

### Rapid 3D imaging of single neurons in the whole brain at a time scale of minutes

Our geometry with 1-5 μm thin tunable Bessel plane illumination and 1.26-12.6× zoomable detection readily enables isotropic brain (or other organs) imaging at different resolution scales (**Supplementary Fig. 11**). More importantly, the expanded SPB made possible by our CACS computation allows whole-brain (or other organs) imaging with isotropic subcellular resolution and at very high throughput, thereby quickly visualizing massive individual cells with details across the entire brain. In our demonstration of whole-brain imaging of neurons, a cleared mouse brain (Thy1-GFP-M, ∼10 × 8 × 5 mm^3^) was first imaged using the 3.2× Bessel sheet mode (FOV 4.2 mm, **Fig. 3a1**) under two views, each containing six stacks (**Fig. 3a2**). Then, tile stitching^38^ for each view (**Fig. 3b1, 2**) followed by a two-view fusion^39^ was applied to obtain the 3D image of complete brain (**Fig. 3a2, 3b3**; 2 μm voxel). CACS was applied at the last step (**Fig. 3a3**) to generate a final output (0.5 μm voxel) with resolved nerve details comparable to those obtained with the 12.6× Bessel sheet (**Fig. 3b4, c, Supplementary Fig. 11, Video S3**). The 10-minute rapid acquisition combined with parallel computation using multiple GPUs permits the efficient reconstruction of a digital brain with isotropic subcellular-resolution over a 400 mm^3^ volume (**Fig. 3c, Methods, Supplementary Note Table 2, Video S4**). Magnified views of three small regions in cortex, cerebellum, and midbrain by 3.2× Bessel sheet, 12.6× Bessel sheet, and 3.2× Bessel-CACS were also compared (**Fig. 3c**), revealing the various neuron morphology across the large-scale brain. The fine nerve fibers, such as apical dendrites, densely packed in these regions were successfully super-resolved in three dimensions by CACS procedure. Furthermore, through referring to the high-resolution 12.6× Bessel sheet result, the recovery accuracy of CACS at whole-brain scale was also verified by comparing the NRMSE and SSIM values of 12 selected sub-regions (100 × 100 × 100 μm^3^) across the entire reconstructed brain (**Fig. 3d**). Therefore, brain segmentations together with accurate quantifications could be implemented at system level (**Fig. 3a4**). For example, we precisely traced interregional neuron projections, which are important for understanding the functionality of the brain. Compared with the raw image, the CACS visualized more nerve fibers (**Fig. 3c**), thus enabling neuronal trajectories with abundant details to be presented in three dimensions (**Fig. 3d, Supplementary Video S3**). By referring to the standard mouse brain atlas (Allen Brain Institute)^3,40^, anatomical information was annotated onto the CACS Bessel brain Atlas (**Supplementary Video S5**), and the trajectories of four long-distance (LD) projection neurons were traced and registered to the annotations, revealing how their pathways were broadcast across the anatomical regions (**Methods, Supplementary Fig. 12**). With isotropic submicron resolution, 13 densely-packed Pyramidal neurons could be also identified and traced through the entire cortex to the striatum (∼2 × 2 × 2 mm^3^), as shown in **Supplementary Fig. 12** and **Video S6**. This quantitative analysis was implemented in a Thy1-GFP-M mouse with massive numbers of neurons being labelled, and this procedure would be even more efficient if the brain was more specifically labelled (e.g., by virus tracer).

**Fig. 3.**
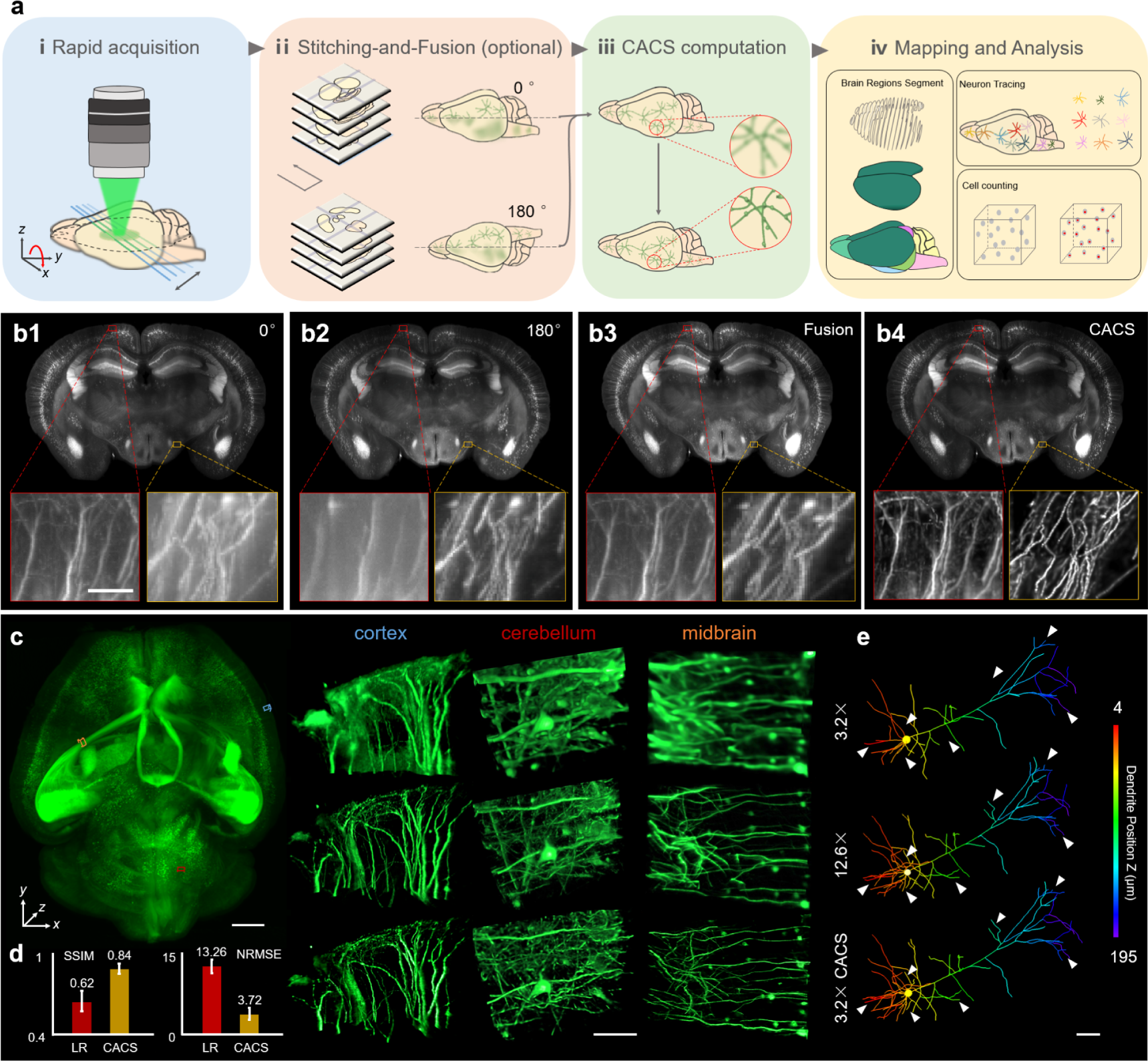
Whole-brain mapping pipeline. **a**, Flow chart of whole-brain data acquisition and processing, which includes: **i**, rapid image acquisition by 3.2× Bessel sheet, with total 12 tiles under 2 views acquired in ∼10 minutes. **ii**, tile stitching for each view followed by 2-view weighted fusion, to reconstruct a scattering-reduced whole brain at low resolution (2-μm voxel). The full implementation of this step would be not necessary if the sample is optimally cleared or imaged under lower magnification. **iii**, CACS computation to recover a digital whole brain with improved resolution (0.5-μm voxel) and SNR. **iv**, quantitative analyses, such as brain region segmentation, neuron tracing and cell counting, based on high-quality whole brain reconstruction. **b**, MIPs of reconstructed coronal plane (***x-z***, 150-μm thickness) in whole brain by 0°, 180° single view Bessel sheet, 2-view fusion and CACS computation. The magnified views of two small regions selected from ***z*** = 100 and ***z*** = 4500 μm depth further show the effect of 2-view fusion and CACS. Scale bar, 50 μm. **c**, 3D visualization of a whole digital mouse brain (left, ∼400 mm3, 3-trillion voxels) reconstructed from the 3.2× Bessel-CACS results, whose raw data was rapidly acquired in ∼10 minutes. Three magnified volumes from the cortex (blue), cerebellum (red) and midbrain (orange) regions by raw 3.2× Bessel sheet (top row), 3.2× Bessel-CACS (middle row), and 12.6× Bessel sheet (bottom row) modes are compared to show the super-resolution capability of CACS procedure. Scale bars, 1 mm for whole brain visualization, 50 μm for ROIs. **d**, SSIM and NRMSE values of 12 brain sub-regions (100 × 100 × 100 μm3) images by raw Bessel sheet and Bessel-CACS. The quantitative analyses verify a high recovery fidelity of CACS results as compared to the high-resolution ground truths. **e**, Tracing of a single pyramidal neuron under three modes. The same neuron imaged by three modes was segmented and traced using Imaris. The results clearly show the improved segmentation accuracy (white arrows) brought by CACS. Scale bar, 50 μm.

### 3D imaging and cell counting in half mouse brain

In addition to the tracing of neuronal projections, the cyto-structures at different brain sub-regions were also explored using CACS Bessel sheet imaging (**Fig. 4**). In order to obtain numbers of cell in different sub-regions with the irregular and diversified morphologies of cells, we labelled nuclei with propidium iodide (PI), an organic small molecule staining DNA. Almost all cell nuclei including motor axons and retinal ganglion cells were stained, causing very dense fluorescing (**Fig. 4a**). Raw 3D images of the nuclei in half brain quickly obtained by 2× CACS sheet mode (**Fig. 4a**; in ∼4 minutes), signals remained highly overlapped and undistinguishable (**Fig. 4b1**). Identically, CACS offered a 4× resolution enhancement in each dimension to recover images with resolution even better than 8× Bessel sheet, to discern single nuclei (**Fig. 4b2, Supplementary Video S7**). The vignette high-resolution views show that cell counting based on the CACS was as accurate as that using an 8× Bessel sheet (**Fig. 4c2, 3, Supplementary Video S8**). After anatomical annotation was applied by registering the image volume to the aforementioned Allen brain atlas (ABA), the labelled cells could be properly segmented (**Fig. 4d-g, Supplementary Video S9**), with each individual cell being assigned an anatomical identity, such as isocortex, hippocampus formation, olfactory area, cerebral nuclei, cortical subplate, thalamus, hypothalamus, midbrain, hindbrain, or cerebellum, as shown in **Fig. 4h, i**. We did not observe morphology- or size-dependent errors in cell detection in different regions. By registering and annotating the detected cells to anatomical areas in the CACS Bessel Atlas, cell number and density information for these brain regions was quantified (**Supplementary Fig. 13, 14 and Video S9**). As shown in **Fig. 4j**, the total cell number in the half brain was ∼3.5×10^7^, and the average density was ∼2.4×10^5^ cells per mm^3^. Amongst the primary brain regions, the Isocortex showed the highest number of cells at 8.6×10^6^, which compares with 2.3×10^6^, 7.3×10^6^, 5.3×10^5^, 3.6×10^6^, 2.4×10^6^, 1.4×10^6^, 2.8×10^6^, 1.3×10^6^ and 3.7×10^6^ cells in the Olfactory area (OLF), Cerebellum (CB), Cortical subplate (CTXsp), Cerebral nuclei (CNU), Hippocampal formation (HPF), Hypothalamus (HY), Midbrain (MB), Thalamus (TH), and Hindbrain (HB), respectively. The cerebellum had the highest cell density of ∼4×10^5^ cells per mm^3^, compared with 2.5×10^5^, 2.4×10^5^, 2.3×10^5^, 2.1×10^5^, 2.1×10^5^, 2.6×10^5^, 2.2×10^5^, 2.0×10^5^ and 1.7×10^5^ in the other regions listed above. Although the current results were obtained using a PI-stained half brain, in which ∼50% of all motor axons and retinal ganglion cells were counted, our results are consistent with previously reported work^2^.

**Fig. 4.**
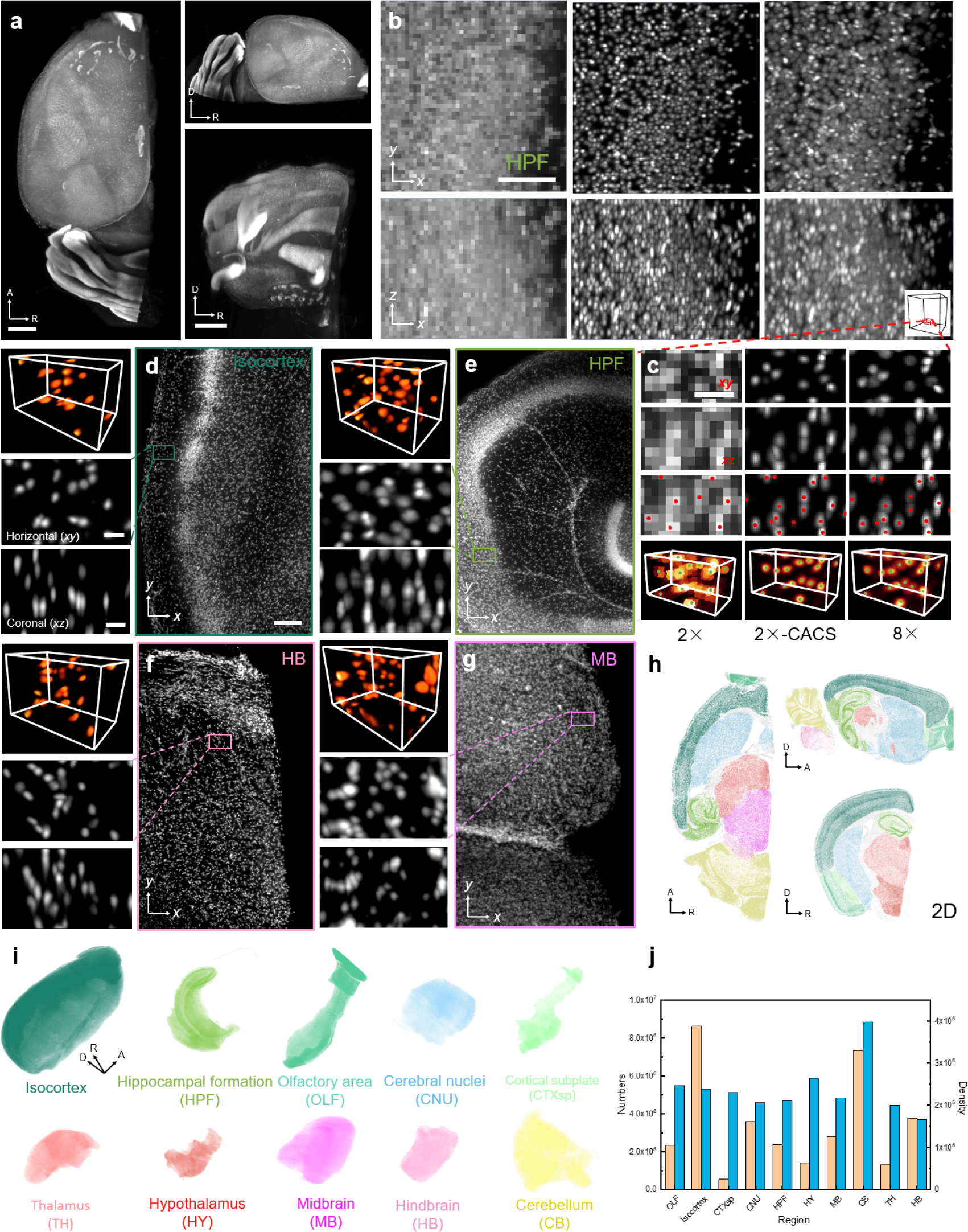
Cell counting of propidium iodide (PI)-labelled half mouse brain. **a**, Coronal view, transverse view and sagittal view of PI-labelled half mouse brain image by 2× CACS Bessel sheet. Scale bar, 1 mm. **b**, A selected ROI in HPF, resolved by 2× Bessel sheet (left column), 2× CACS Bessel sheet (middle column), and 8× Bessel sheet (right column). **c**, Magnified views of a small volume (red box) in **b**, showing the different accuracy of nuclei based identification base on three types of images. Scale bar, 20 μm in **b** and 10 μm in **c. d-g**, 3D visualization of four high-resolution ROIs from Isocortex, HPF, HB and MB regions in the reconstructed half brain. Scale bar, 50 μm in large box and 10 μm in small box. The results clearly show single-cell resolution in three dimensions, which is sufficient for cell counting. **h**, 10 manually segmented brain regions in the half brain, shown in transverse, coronal, and sagittal views. **i**, 3D visualization of the 10 sub-regions indicated by different colors. **j**, Calculated nuclei number and density of each sub-region in half brain.

### Dual-color 3D imaging and quantitative analyses of neuro-muscular junctions

We further demonstrated dual-color CACS Bessel sheet imaging of motor endplate (MEP, α-BTX) and peripheral nerves (Thy1-YFP) in mouse muscles. The overall spatial distribution of MEPs in the muscle tissue and the detailed neuromuscular junction (NMJ) occupancy defined as the volume ratio of the underlying postsynaptic MEPs (red) occupied by presynaptic vesicles (green), were both visualized and analyzed using the CACS Bessel sheet microscopy. We first finished high-throughput screening (3.2× Bessel sheet) of two gastrocnemius muscles and two tibialis anterior muscles (∼720 mm^3^ in total) rapidly in ∼20 minutes (**Fig. 5a-d, Supplementary Video S10**). The image-based segmentation of the MEPs and nerves across whole-muscle scale was then enabled as a result of the quick 3D visualization (insets in **a-d, Supplementary Video S11**). Image results of two ROIs in gastrocnemius (**Fig. 5a)** by raw 3.2× Bessel sheet, 3.2× Bessel-CACS, and high-resolution 12.6× Bessel sheet are compared in **Fig. 5e**, to show the quality of 3.2× Bessel-CACS results being as high as those from 12.6× results. The magnified views of single NMJs (six white boxes) further show that CACS could reveal fine neuromuscular structures in single NMJs (**Supplementary Video S11**). The number of MEPs and their density in each muscle were thereby quantified based on the large-scale 3D images (**Fig. 5f**). We found that although the total MEP number varied in each muscle, their density remained relatively close. Furthermore, owing to the high spatial resolution achieved across whole-muscle scale, our method also permitted the accurate measurement of presynaptic and postsynaptic structural volumes in single NMJs and the consequent calculation of NMJ occupancy (right, **Fig. 5g**), which are not possible for conventional 3.2× Bessel results (left, **Fig. 5g**). These complete visualization and quantitative analyses enabled by CACS Bessel sheet imaging indicate both the contraction capability of muscle tissues at a macro scale, and the delivery efficiency of neurotransmitters at nerve endings on a micro-scale, thereby potentially benefiting the comprehensive understanding of neuromuscular functional activities and improving the diagnosis and development of therapy for related diseases^41,42^.

**Fig. 5.**
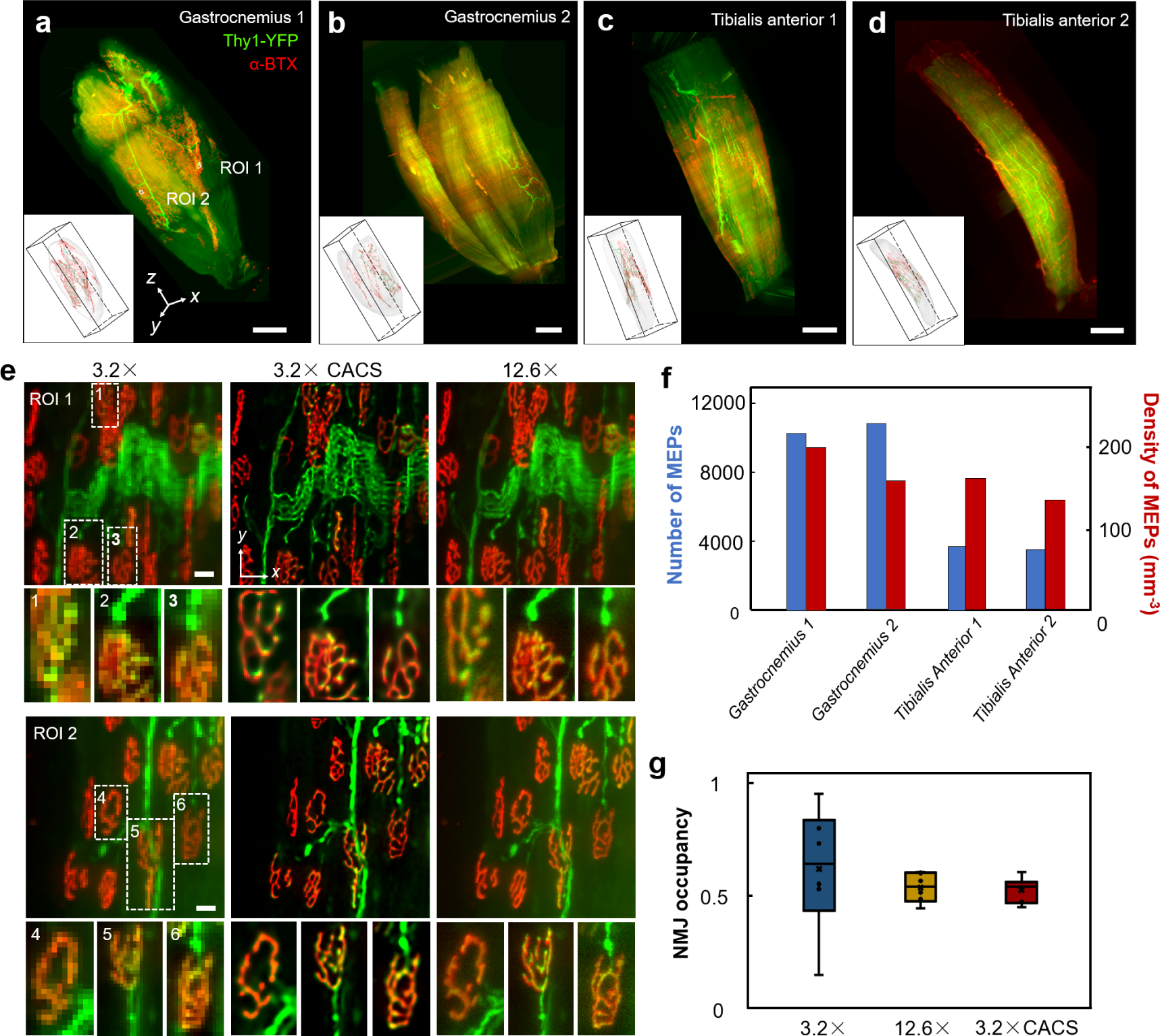
Dual-color CACS imaging and quantitative analyses of neuromuscular junctions in mouse Gastrocnemius and Tibialis anterior. **a-d**, Dual-color, 3D visualization of Gastrocnemius and Tibialis anteriors, showing both the presynaptic terminal bouton (green, Thy1-YFP) and postsynaptic acetylcholine receptors (red, α-BTX). Insets show image-based segmentation of presynaptic terminal and MEPs. **e**, Comparison of two ROIs (100 × 100 × 100 μm3) in **a**, by 3.2× (left column), 3.2× CACS (middle column) and 12.6× Bessel sheet (right column). Six magnified views (white boxes, 1, 2, 3 for ROI1 and 4, 5, 6 for ROI2) further demonstrate the structural information of single NMJs. Scale bar, 10 μm. **f**, Macro-scale quantitative analysis of MEP, including total numbers and densities of MEPs, in two types of muscles. **g**, Neuromuscular junction (NMJ) occupancy calculated based on abovementioned 3.2×, 3.2× CACS and 12.6× Bessel sheet images that show different level of resolution. The result from raw 3.2× image shows an unacceptably large error range, owing to the ambiguous cellular resolution. Here NMJ occupancy was calculated as (presynaptic volume/postsynaptic volume) ×100%.

## Discussion

In summary, the marriage of the CACS strategy with long-range Bessel plane illumination enables quick 3D imaging of millimeter-to-centimeter sized samples in seconds to minutes, with subcellular isotropic resolution and high throughput of up to 10 gigavoxels per second. As the size of optically-cleared and labelled sample becomes larger and larger nowadays, such technical advances of resolution/throughput shown in our method also becomes increasingly valuable, especially for the *intoto* mapping of entire organs/organisms at variable scales. Highly inclined LSM has been used for the 3D anatomical mapping of thick organs where autofluorescence and out-of-focus excitation would otherwise be prohibitive under wide-field illumination. With the thinner light-sheets of the Bessel beam, the effective FOV would be less compromised, and axial resolution would be further improved. The introduction of dual-side illumination and dual-view detection would further reduce light depletion in thick tissue and permit the complete interrogation of entire organs. As currently configured, the key step to maintaining thin and wide Bessel plane illumination under a large FOV is to synchronize the scanned beam with the confocal electronic slit of the camera, to eliminate excitation from the side-lobes. This could possibly be improved by direct generation of a side-lobe-reduced Bessel sheet using recent masking techniques^43^. At the other extreme, the possible introduction of two photon excitation may be furthermore suited for improved depth penetration when imaging more scattering samples.

Despite the use of plane illumination, current microscopes have throughput limited to megavoxels per second. This causes issues in 3D anatomical mapping, in which trillion-voxel data is frequently required. Therefore, the long-time acquisition required for mechanical image stitching prevents a variety of histological, pathological, and neuroanatomical research being implemented on a large scale. Our CACS procedure uses a single image stack to three-dimensionally improve the contrast and resolution limited by the pixilation and numerical aperture, computationally transforming our Bessel microscope into a gigavoxel-throughput-boosted imager that can achieve isotropic 3D super-resolution imaging of mesoscale samples without the pain of a long acquisition lasting for hours to days. Considering that signal characteristics such as sparsity and degree of disorder vary drastically in large specimens, it is difficult for regular CS computation to process the signals appropriately, because it is uncertain what regularization factor should be applied. Our introduction of a content-aware feedback strategy can reasonably extract the signal characteristics and thereby calculate the optimal regularization parameter for each image block automatically, allowing recovery of signals on a large scale with minimal artefacts, as well as obvious resolution improvement, which is unmet by conventional CS (**Supplementary Note Fig. 1**). Our 3D CACS computation is unsupervised, without the need for prior knowledge from a higher resolution database, and is highly parallelized for GPU acceleration. It should be noted that CS processing is efficient for the recovery of under-sampled signals, a condition particularly satisfied in our low-magnification large-FOV imaging setup. Moreover, although compressed sensing is known to have certain limitations for processing signals containing complicated structural information^44,45^, its effect on line-like neurons and point-like nuclei was notably improved after the content-aware adaption to the signal characteristics involved. We expect our method to potentially bring new insights for computational imaging techniques, so that we can keep pushing the optical throughput limit to extract ever more spatiotemporal information from biological specimens. We also believe that combinations with other computational techniques, such as neural network-enabled data training, could further increase the ability of our method to achieve more robust and versatile recovery of various complicated signals in different types of specimens.

## Acknowledgements

We thank Shangbang Gao, Luoying Zhang, Haohong Li, Man Jiang, Bo Xiong for discussions and comments on the manuscript, Yunyun Han for the discussion on the imaging and brain visualization, Zhilong Yu and Hao Zhang for the support on the code implementation. This work was supported by the National Key R&D program of China (2017YFA0700501, D.Z. and P.F.), the National Natural Science Foundation of China (21874052 for P.F., 61860206009 for D.Z., 81873793 for W.M.), the Innovation Fund of WNLO (P.F. and D.Z.) and the Junior Thousand Talents Program of China (P.F.).

## Author contributions

P.F. conceived the idea. P.F., D.Z. and W.M. oversaw the project. C.F. and X.W. developed the optical setups. C.F. developed the programs. C.F., T.C., T.Y., Y.H., W.F., P.W., and Y.L. conducted the experiments, processed the images, and visualized the data. C.F., T.Y., D.Z., W.M. and P.F. analyzed the data and wrote the paper.

## Methods

### Zoomable Bessel plane illumination microscopy setup

The optical setups of zoomable Bessel plane illumination microscope were detailed in **Supplementary Figs. 1, 2, Video S1**. Our Bessel plane illumination microscope (**Supplementary Fig. 1, Video S1**) sweeps a long circular Bessel beam in the ***y*** direction across the focal plane of the detection objective to create a large-scale scanned light-sheet and yield an image at a single ***z***-plane within the specimen. To maintain sufficient laser intensity at large scale, we used a large apex axicon (176e, AX252-A, Thorlabs) instead of mask to create annular beam at a plane conjugate to both the galvanometer and the rear pupil of the excitation objective. For the Bessel beam, its axial extent of central maximum (beam thickness) and diffraction distance (beam length) are proportional to the magnification factor of the illumination objective and its square, respectively. To suppress the laser attenuation/scattering from samples, we introduced two opposite plane illumination sources from dual sides of the brain. A macro-view microscope (Olympus MVX10, 0.63× to 6.3×, combined with a 2× detection objective lens MV PLAPO 2XC) providing zoomable magnification from 1.26×/0.14 to 12.6×/0.5, together with an sCMOS camera (Hamamatsu ORCA-Flash4.0 V2) were used as the fluorescence detection unit, providing a tunable FOV from 1 to 10 mm and effective lateral resolution from ∼1 μm to 10 μm for large specimens. To achieve isotropic resolution, we used a set of switchable long working distance objectives (Mitutoyo, 2×/0.055, or 5×/0.14, or 10×/0.28) from each side to form a tunable dual-side illuminating Bessel beam with axial extent and diffraction length matched with the varying lateral resolutions and FOV, respectively. To achieve side-lobe free imaging based on electronic confocal slit, we formed a linear array which contains 2-4 rows of active pixels sweeping at the sensor plane along ***y*** direction, and tightly synchronized its movement with the central maximum of scanned Bessel beam (**Supplementary Video S1**). During image acquisition, the 2D galvo mirror (GVS012, Thorlabs) that scanned the Bessel beam into plane, the sCMOS camera that continuously recorded the fluorescence images at high speed up to ∼50 frames per second, the motorized ***xyz*** stage (Z825B, Thorlabs combined with SST59D3306, Huatian) that moved the sample in three dimensions, and the motorized flip mirror that switched the illumination between dual-side beams, were all controlled by analog signals generated by a data acquisition device (PCIe-6259, National Instruments). To enable high-throughput volumetric imaging of specimen, a customized sample holder rapidly moved the sample across the light-sheet along the ***z*** direction, with plane images at different depths consecutively obtained (**Supplementary Fig. 2**). The holder can also flip the sample through 180° to allow imaging under two views, so that the acquired dual-view images can be fused to further suppress fluorescence depletion from deep tissue. For the imaging of an entire organ, the sample was flipped 180 degrees up-side down to be imaged twice. Due to the corrosive reactive index matching immersion solution, a micro-motor was sealed inside a chemical-resistant resin box and attached to the sample holder to fulfill such precise rotation. The two-view images were then registered and fused using multi-view fusion method^39^. A LabVIEW (National Instruments) program was developed to synchronize these parts, to automatically realize the line-synchronized scanning, ***z***-scanning, tile stitching, and sample rotation in order.

### CACS computation procedure

If the signals are not dense and incoherent at the same time, CS allows the recovery of them from incomplete measurement, which is our case in large-FOV Bessel sheet imaging. To correlate the high-resolution image to be recovered with the real measurement, we first used a 3D Gaussian-Bessel function as PSF to characterize the unit response of the Bessel sheet system. The lateral and axial extents of the PSF are determined by a variety of imaging parameters, such as NA of illumination and detection objectives, and camera’s spatial sampling rate. Then a measurement matrix ***A*** could be generated by Fourier transformation of the synthetic PSF. Here ***A*** represents the signal degradation operation including the optical blurring by system optics and down-sampling by the camera digitalization in Fourier space, transforming the high-resolution target into low-resolution measurement (**Supplementary Video S2**). Then the Fourier expression of high-resolution image ***x*** can be recovered from the Fourier expression of low-resolution measurement ***y*** by iteratively solving following equation in Fourier domain using a steepest descent method:

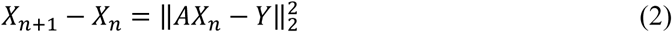

The recovered image can be finally obtained via transforming the solved ***x*** back into spatial domain. As we have known, compressed sensing could show very different effects on signals with different labelling density, which is particularly common to large samples. An ***L1***-normalization term was thus introduced in our method to regularize the equation as following:

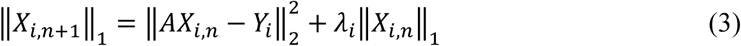

Here the weighting factor ***λ***_***i***_ substantially balances the image results from more like the original measurement to more like the CS recovery. The value of ***λ***_***i***_ is calculated by the multiply of ***α***_***i***_ and ***β***_***i***_, where ***α***_***i***_ is an index indicating the signal density of input image (**Supplementary Note 3**), and ***β***_***i***_ is inversely proportional to the entropy of the input image (**Supplementary Note 3**). Therefore, the choice of ***λ***_***i***_ is highly adapted to the signals characteristics which could vary a lot at large scale, our content-aware processing procedure divides the raw image, e.g, half brain image with dimension 10 mm × 4 mm × 5 mm, into a series of small volumes ***x***_***i***_, e.g., blocks with 100 × 100 × 100 pixels in which the signals can be considered as uniformly distributed, and then calculates the regularization factor for each volume to obtain the optimal CS result ***x***_***i***_. Multiple ***x***_***i***_ are transformed back to high-resolution image blocks, which would be stitched into final output (**Supplementary Video S2**). It should be noted that the CACS computation can be highly parallelized by using GPU acceleration.

### Rapid 3D isotropic imaging of mouse brain at various scales

The improved axial resolution obtained from the tunable Bessel plane illumination and zoomable detection enables easy brain (or other organs) imaging at isotropic resolution and various scales. As a reference, four clarified mouse brains (Tg: Thy1-GFP-M) were quickly screened at a throughput of one brain min^-1^ and ∼10 μm isotropic resolution (**Supplementary Fig. 11b)**. Each brain (10 mm × 8 mm × 5 mm) was imaged under the lowest 1.26× with our MVX10 microscope. Two 2×/0.055 excitation objectives generated a Bessel light-sheet with a thickness of 5 μm from dual-side. Considering ultra-high resolution in fast screening mode was not required, we set the scanning step size as 5 μm (voxel size 5.16 μm × 5.16 μm × 5 μm), to final achieve ∼10 μm isotropic resolution. Under the 1.26× dual-side Bessel sheet mode, we acquired a 3D image stack containing ∼600 whole-brain slices at each side in each view, finally obtaining four image stacks containing over 2000 large-FOV image slices for both sides of both views in ∼60 seconds. These four sets of image stacks were then computationally stitched and fused together, forming a final 3D image of the whole brain in a few minutes^2^. By obtaining the coarse 3D structures of the whole brains using the widest Bessel sheet (**Supplementary Fig. 11b**), the transverse (***x-y***), coronal (***x-z***), and sagittal (***y-z***) views could be extracted from the reconstructed brains to quickly identify brain regions where desired signals were present (**Supplementary Fig. 11c**). Then, higher-resolution imaging of any region of interest was possible using the 12.6× Bessel sheet mode. For example, No. 2 brain with showing the best signal distributions was chosen after the fast screening. We then imaged three ∼1 mm^3^ regions of interest (ROIs) in the cortex, hippocampus, and cerebellum of No. 2 brain (**Supplementary Fig. 11b**) at an imaging speed of ∼0.01 mm^3^ s^-1^ and an isotropic resolution of ∼1.5 μm, respectively. For ROIs imaging under 12.6× magnification, we switched the motorized flip mirror (MFF101/M, Thorlabs) to choose single side illumination (left or right), depending on the position of ROIs. The scanning step size was 0.5 μm for sufficient ***z*** sampling. Therefore, the voxel size was 0.516 μm × 0.516 μm × 0.5 μm. Vignette high-resolution views of these volumes are shown in **Supplementary Fig. 11d-f**, detailing the various neuron morphologies across the brain. Besides the identification of different types of neuron cell bodies (e.g., pyramid neurons in **Supplementary Fig. 11e**, and astrocytes in **Supplementary Fig. 11f**), nerve fibers such as densely packed apical dendrites (**Supplementary Fig. 11d**) were also clearly resolved in three dimensions.

### Whole-brain CACS imaging procedure

For high-resolution mapping of whole mouse brain to enable single-neuron-level quantitative analysis across whole brain, the choice of repetitive tile stitching, e.g., around 100 times under 12.6× magnification, is not only very time consuming, estimated with over 15 hours, but also brings substantial photo-bleaching. Instead, we imaged the whole brain at lower 3.2× Bessel sheet mode and used CACS to computationally improve the compromised resolution. With 4.16 by 4.16 mm FOV under 3.2×, we sequentially obtained 6 volumes of the different regions of the brain with each one containing 1500 frames for 3 mm ***z***-depth (2 μm step-size), and stitched them into a whole 3D image. Then, two views (0° and 180°) of stitched images were iteratively registered for reconstructing a whole brain with exhibiting complete structural information while presenting spatial resolution ∼4 μm (2 μm voxel), which remains insufficient for single-cell/ neuron analysis. Nevertheless, CACS was applied with automatically subdividing the whole brain into thousands of blocks for being processed using 4 GPUs (RTX2080ti) in parallel block by block (100 × 100 × 100 voxels for each block), owing to the memory limit of the GPUs. Finally, the super-resolved patches were automatically stitched together into a complete 3.5 trillion-voxel image (0.5-μm iso-voxel size) with an acquisition time ∼10 minutes, which is compared to tens of hours by other methods^1^, and computation time ∼5 hours which is estimated similar with the time consumption for stitching and fusion under 12.6×.

It should be noted that the combination of Bessel sheet with CACS is very flexible, providing throughput improvement for zoomable imaging from 1.26× to 12.6×. It is also easy to modify the LabVIEW program to conduct imaging under different modes/magnifications. Thus, it’s beyond the imaging of brain we demonstrate, being widely suited for anatomical imaging of various large samples, such as mouse muscles shown in **Fig. 5**.

### Dual-channel mouse muscle tissues imaging procedure

For dual-color imaging of gastrocnemius muscles and tibialis anterior muscles, we excited neurons and presynaptic membranes with a 488 nm laser and postsynaptic membranes (also known as MEPs) with a 637 nm laser (RGB-637/532/488-60mW, Changchun New Industries Optoelectronics Technology Co., Ltd.). Under 3.2× Bessel sheet mode, we stitched twice and three times for entire tibialis anterior muscle and gastrocnemius muscle, respectively. The thickness of all the muscles were around 3 mm, hence the imaging under 2 views for multiview-fusion was not necessary. It merely took around 5 minutes to complete dual-channel imaging of each muscle, and the total acquisition time was approximately 20 minutes.

### Sample preparation

uDISCO^19^ clearing method was used to clarify the dissected wild-type mouse brain with cell nuclei stained by PI dye, neuron-labelled mouse brain blocks (Tg: Thy1-GFP-M). FDISCO clearing method^46^ was used to clarify the neuron-labelled mouse muscles (Tg: Thy1-YFP-16) with α-BTX tagging the motor endplate. PEGASOS clearing method^47^ was used to clear the Thy1-GFP-M whole mouse brain, The cleared organ samples were harden and thus could be firmly mounted onto our customized holders (**Supplementary Fig. 2**). The BABB-D4 for uDISCO, DBE for FDISCO or BB-PEG for PEGASOS immersion liquid was filled in the sample chamber for refractive index-matched imaging with least aberration. For measuring the system’s point spread function, fluorescent beads (Lumisphere, 1%w/v, 0.5 μm, Polystyrene) were embedded into a specifically formulated resin (DER332, DER736, IPDA, 12: 3.8: 2.7), the refractive index of which was equal to BABB-D4, to form a rigid sample that can be clamped by the holder and imaged in the chamber.

### Whole-brain visualization and brain region registration to ABA

After the CACS enhancement on the low-resolution whole brain data, we used an adaptive registration method to three-dimensionally map the brain to the standard Allen Brain Atlas. Since the Allen Brain Atlas was reconstituted from a series of coronal-slice images, firstly we also re-orientated our reconstructed image from horizontal view to coronal view and automatically pre-aligned to the ABA using Elastix. This pre-aligned brain was then resized into low-resolution and high-resolution groups. Next, we finely registered the low-resolution group to the ABA and obtained the transform correspondence, which was then applied to the high-resolution group to obtain a registered and transformed high-resolution brain. This mapped brain atlas was finally visualized in Imaris to facilitate the neuron analysis. Then a manual adjustment was followed to improve the accuracy. Finally, this mapped brain atlas was visualized in Imaris to facilitate the neuron and cell number analysis. With the creation of the atlas, the neurons or cell nuclei localized to different encephalic regions (such as isocortex, hippocampus, cerebellum and midbrain) could be identified (**Fig. 4i**), and *intoto* mapped out at a whole-brain scale (**Fig. 4h, Supplementary Video S9**). Then, the neuron population and the cell nuclei density in different encephalic regions were quantified by calculating the volume of the regions and counting the identified cell bodies within them (**Fig. 4j, Supplementary Video S9**).

### Neuron tracing

The whole brain data was down-sampled 2 times and segmented manually using the commercial Imaris software. With registering our brain to ABA, we obtained the anatomical annotation for all the segmented areas. The Autopath Mode of the Filament module was applied to trace long-distance neurons. We first assigned one point on a long-distance neuron to initiate the tracing. Then, Imaris automatically calculated the pathway in accordance with image data, reconstructed the 3D morphology and linked it with the previous part. This automatic procedure would repeat several times until the whole neuron, which could also be recognized by human’s eye, was reconstructed.

### Cell counting

The Spots module and Surface module of commercial Imaris software was used to count cells in various anatomical regions of CACS-reconstructed half brain (full resolution, ∼400 gigavoxels). We first separated several primary brain regions into different channels in Surface module. Then automatic creation in Spots module was applied to count cells number for each single channel which represents an encephalic region. To achieve accurate counting, the essential parts were the appropriate estimate of cell bodies’ diameter and filtration of the chosen cells by tuning the quality parameters. This automatic counting procedure was also aided by human’s crosscheck, which herein severed as golden standard. After obtaining the total number of cells in each primary brain region, according to the sub-region ranges divided by the Surface module, Imaris could also calculate the volume of each primary brain region. Then, with knowing the number of cell nuclei and the volume of each segmented brain region, the density of the cell nuclei inside each brain region could be calculated.

### Counting of MEPs and calculation of NMJ occupancy

The method to count MEPs was similar to the aforementioned cell nuclei numbers counting with PI-labelled mouse half brain. Under the low-resolution 3.2× magnification, the optical blur brought by the low NA and low sampling rate caused the details inside a single MEP to be lost, and the entire MEP blurred to be a point, which in turn helped us use Imaris’ Spot module to count the number. Considering that we only needed the total number and overall density inside the entire muscle, segmentation was no longer necessary. Select the entire region contained in the image and set the threshold according to the diameter of the single MEP, then we could get the total number of MEPs within a single muscle tissue. To calculate the overall density, we needed to calculate the volume of the whole muscle by the method mentioned before.

Calculating NMJ occupancy required accurate measurement of presynaptic and postsynaptic structural volumes in a single NMJ. After importing the dual-channel images (YFP and α-BTX) into Imaris, we randomly selected 10 NMJs for analysis. For each NMJ, we used the Surface module to calculate the volume of the presynaptic and postsynaptic structural volumes acquired by different channels. The two volume results were divided to obtain the occupancy of a single NMJ.

### Code availability

Custom codes for CACS computation implemented in current study are available from the corresponding authors upon request.

### Data availability

The datasets generated and analyzed in this study are available from the corresponding authors upon request.

