## Supplementary figures and notes for "Minutes-timescale 3D isotropic imaging of entire organs at subcellular resolution by content-aware compressed-sensing light-sheet microscopy"

### Supplementary Information

**Supplementary Figure 1** | Dual-side, dual-mode light-sheet microscopy setup
**Supplementary Figure 2** | Photograph of the built-up system
**Supplementary Figure 3** | Comparison of different plane illumination modes
**Supplementary Figure 4** | Necessity for line synchronization under different magnification/NA setups
**Supplementary Figure 5** | Timing diagram of for line synchronization and unsynchronized dithering modes
**Supplementary Figure 6** | CACS computation for line-like neuron signals at different density
**Supplementary Figure 7** | CACS computation for point-like cell nuclei at different density
**Supplementary Figure 8** | CACS computation for 12.6× finer neuronal sub-structures
**Supplementary Figure 9** | Signal-to-noise-ratio (SNR) comparison of different modes
**Supplementary Figure 10** | Photobleaching rate comparisons of different modes
**Supplementary Figure 11** | Scalable isotropic imaging of neurons in mouse brain
**Supplementary Figure 12** | Tracing dense long-distance projection neurons in 8-week Thy1-GFP-M mouse brain
**Supplementary Figure 13** | Accuracy of compressed sensing in PI-labelled brain
**Supplementary Figure 14** | Segmentation and cell counting for PI-labelled half brain imaged by 2× CACS Bessel sheet
**Supplementary Note 1** | Imaging speed, photobleaching rate and SNR
**Supplementary Note 2** | Image stitching and dual-view image fusion
**Supplementary Note 3** | Content aware regularization in CS
**Supplementary Note Figure 1** | Content-aware calculation of regularization factor
**Supplementary Note Table 1 |** Comparison of different imaging modes
**Supplementary Note Table 2 |** Whole brain imaging with different magnification
**Supplementary Video S1** | Confocally-scanned Bessel light-sheet microscopy
**Supplementary Video S2** | Content aware compressed sensing (CACS) procedure
**Supplementary Video S3 |** Super-resolution of line-like neuron fibers by CACS
**Supplementary Video S4** | 3D imaging of whole mouse brain (neuron tagged) by CACS Bessel sheet microscopy
**Supplementary Video S5** | Whole-brain 3D visualization and segmentation
**Supplementary Video S6** | Tracing long-distance neuronal projections across the entire brain
**Supplementary Video S7 |** Super-resolution of point-like cell nuclei by CACS
**Supplementary Video S8** | 3D imaging of half mouse brain (nuclei stained) by CACS Bessel sheet microscopy
**Supplementary Video S9** | Region-specific cell counting in half mouse brain
**Supplementary Video S10 |** Dual-color 3D imaging of mouse gastrocnemius and tibialis muscles by CACS Bessel sheet microscopy
**Supplementary Video S11 |** Neuron tracing and MEP counting in gastrocnemius muscle

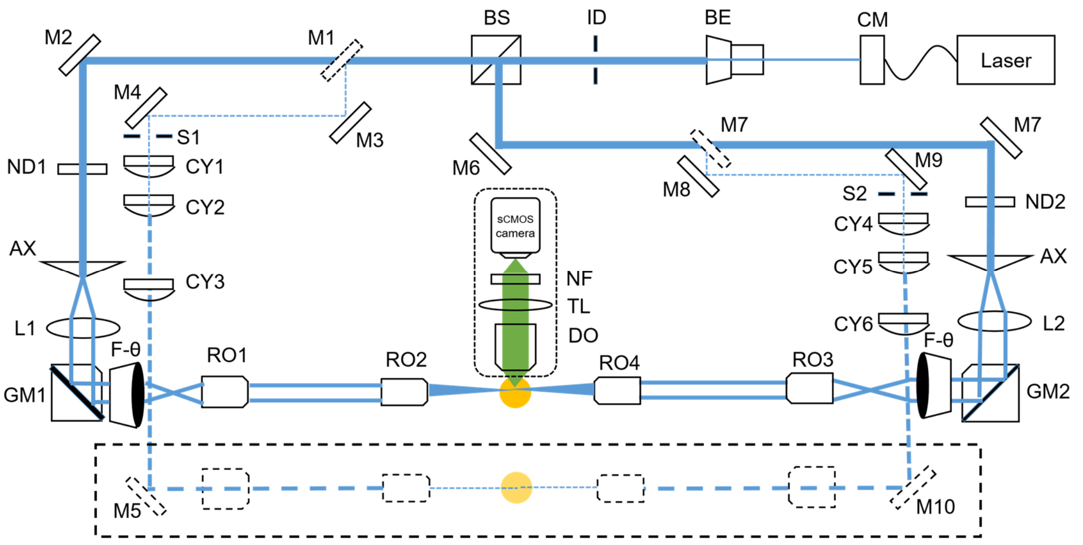


| | |
|---|---|
| CM : Collimator | AX : Axicon |
| BE : Beam Expander | L : Lens |
| S : Slit | F-θ : F-θ lens |
| BS : Beam Splitter | RO : Relay Objective lens |
| M : Mirror | TL : Tube lens |
| ND : Neutral Density Filter | NF : Notch Filter |
| CY : Cylindrical mirror | GM : Galvo Mirror |
| DO : Detective Objective lens | ID : Aperture diaphragm |

**Supplementary Fig. 1 | Dual-side, dual-mode light-sheet setup.** A multi-wavelength laser was collimated and expanded to generate a Gaussian beam with 10 mm diameter. For each side, an axicon (AX) was used to generate a Bessel beam, which was further scanned into a plane by a galvo scanner (GM) and projected onto the sample using two groups of relay lenses (L1, L2, F-theta lenses, RO1-4). Dual Bessel sheets were precisely aligned by finely tuning the *y*-mirror of GM2. Finally, a Bessel plane illumination with widely tunable geometry (1-5 µm thickness, 1-20 mm width, 1-10 mm height) was formed for rapid and high-axial-resolution imaging of large specimen. In our setup, a dual-side Gaussian light sheet was also reserved for quick sample screening as well as being compared to Bessel sheet.

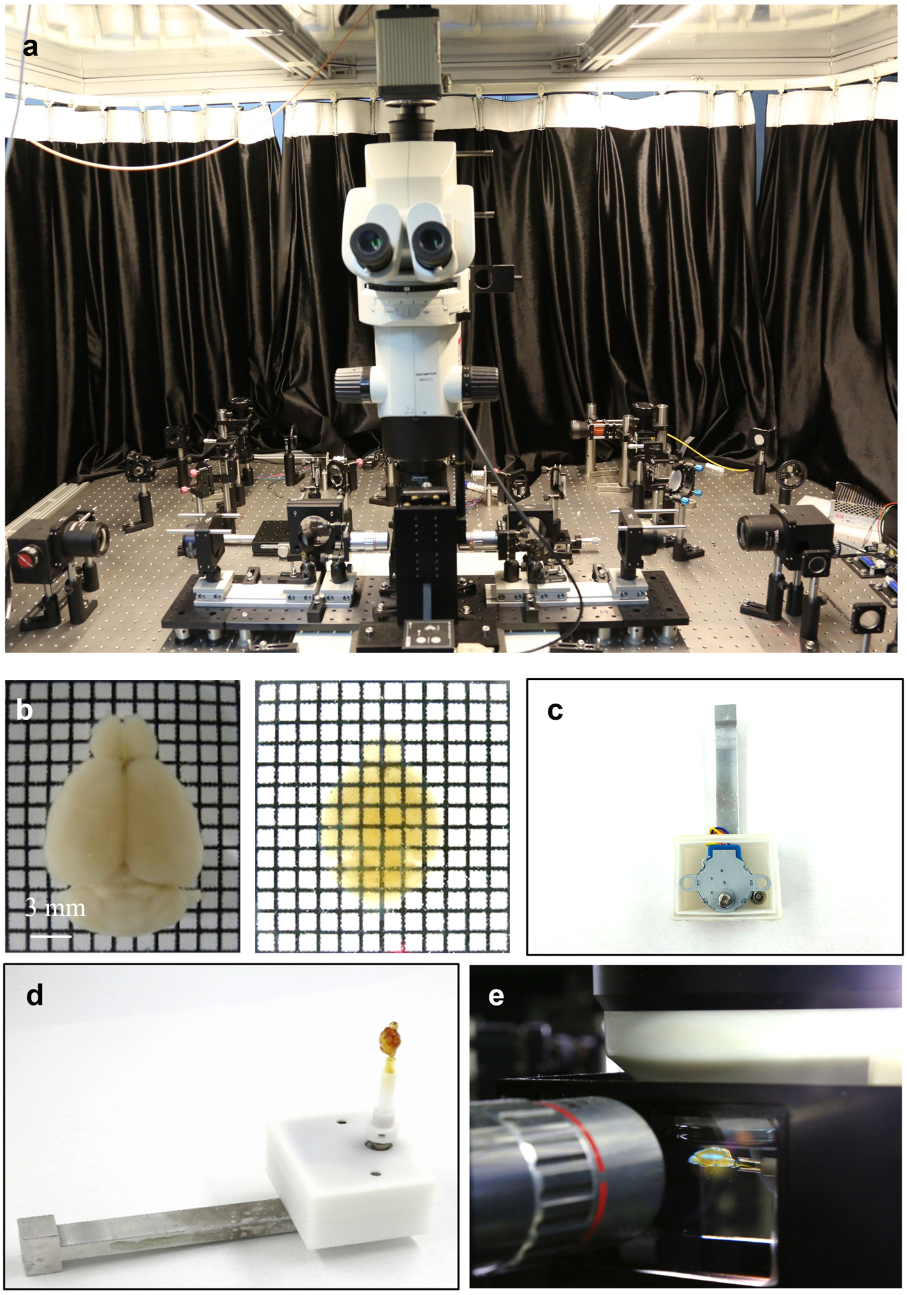


**Supplementary Fig. 2 | The real built-up system**. **a**, The entire view of the microscope constructed on optical bench. **b**, The photographs of an excised whole mouse brain before/after tissue clearing. **c**, A rod-like holder attaching the sample to a 3D translation stage and dipping the sample into a solution chamber. The sample can be three-dimensionally moved and rotated in the chamber for imaging. **d**, A cleared mouse brain sample connected to a waterproof motorized rotator (white box). **e**, Scanned Bessel sheet illuminating a transverse plane of mouse brain.

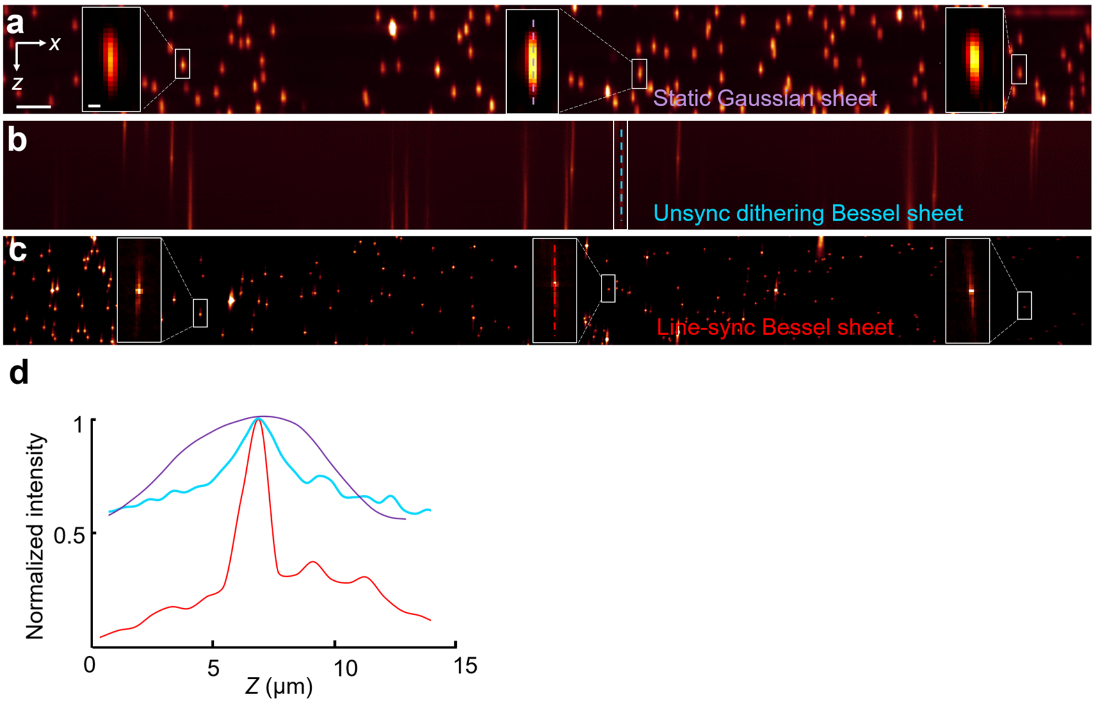


**Supplementary Fig. 3 | Comparison of different plane illumination modes.** We used electronic confocal slit to eliminate side lobe excitation from Bessel beam and thereby increase axial resolution. Under 3.2× detection magnification, with using a 15 μm-thick regular Gaussian sheet as comparison, unsynchronized (high-speed dithering) Bessel sheet caused substantively elongated PSF due to the accumulated axial excitation by side lobes. In contrast, synchronizing scanned Bessel sheet with electronic slit significantly reduced this side effect, yielding near-isotropic PSF with axial extent much shorter than that of Gaussian sheet. **a**-**c**, Axial PSFs measured by Gaussian sheet, unsynchronized Bessel sheet, and synchronized Bessel sheet modes, as shown in **a**-**c**, respectively. **d**, Axial line profiles of the beads resolved by three methods. Scale bar, 50 μm (inset, 5 μm).

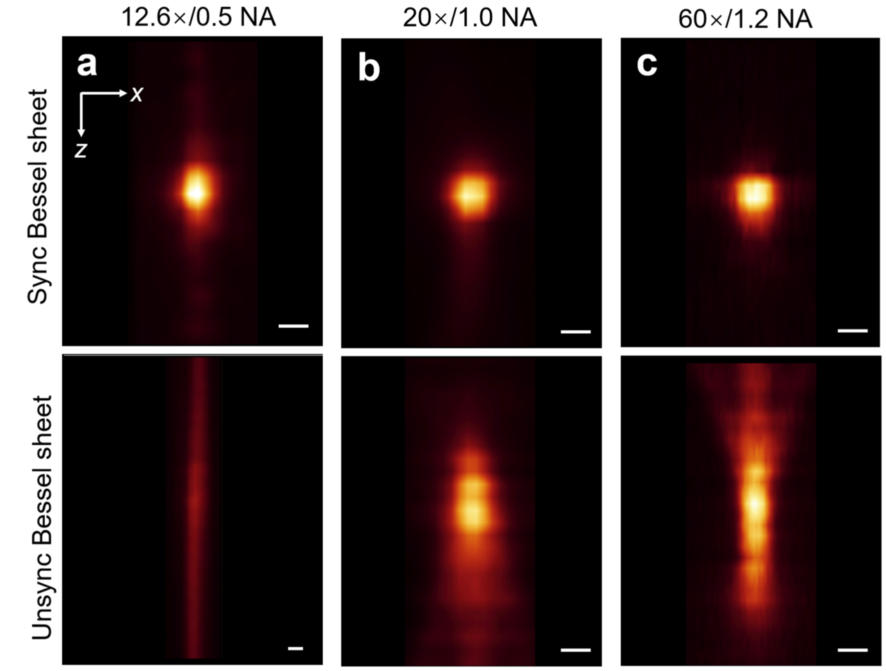


**Supplementary Fig. 4 | Necessity for line synchronization under different magnification/NA setups**. It should be noted that the fluorescence contamination from side lobes' excitation is especially severe under low (3.2×)-to-middle (12.6×) magnification, which is our case, due to the extended depth of focus. PSFs measured by 12.6× Bessel sheet under both synchronized and unsynchronized modes, were compared to the corresponding results from 20×/1.0 NA and 60×/1.2 NA measurements that used the same Bessel sheet illumination (1 μm). With applying line synchronization, all three setups could provide near-isotropic PSFs with similar axial extents. On the other hand, though the PSFs deteriorated in all three setups once synchronization was not included, 12.6×/0.5 setup was obviously more vulnerable to this side effect than 20×/1.0 and 60×/1.2 setups did, owing to its larger depth of focus that received excessive axial signals excited by the side lobes. Therefore, the line synchronization is particularly necessary for our implementation. **a**-**c**, The PSFs measured by 12.6×/0.5, 20×/1.0 and 60×/1.2 setups, respectively, with top row and bottom row showing the synchronized and unsynchronized results, respectively. Scale bar, 0.5 μm.

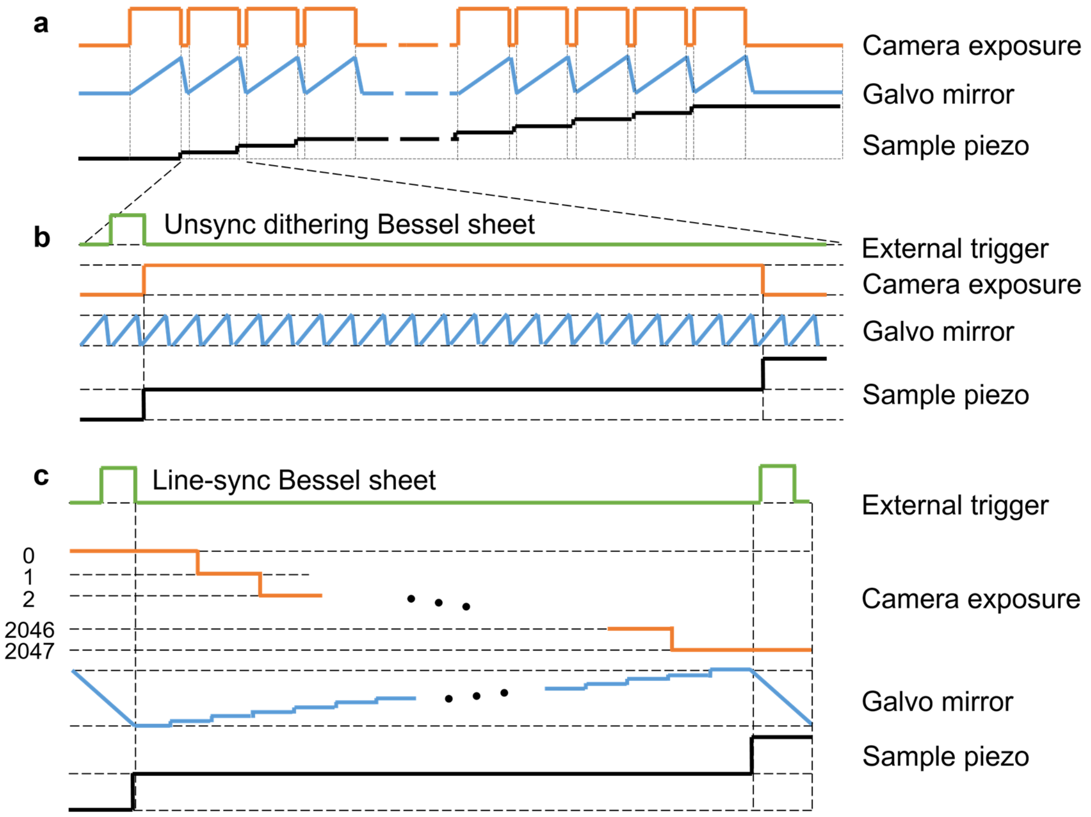


**Supplementary Fig. 5 | Timing diagram of for line synchronization and unsynchronized dithering modes. a**, Control signals for sequential multi-plane imaging. **b**, Control signals for unsynchronized Bessel sheet mode (high-speed dithering) at each plane. In this mode, the camera sensor stayed at global shutter while the beam was scanned back and forth at the sample plane with period much shorter than the exposure time. **c**, Control signals for synchronized Bessel sheet mode. In this mode, the camera sensor generated a rolling exposure line synchronized with the scanned central maximum of Bessel beam. When the beam was scanned from the top to the bottom of sample plane, the narrow active pixel line was simultaneously triggered for rolling with the same velocity and direction, so that the fluorescence signals excited by the central peak of Bessel beam could be always detected while those from the side lobes were always blocked.

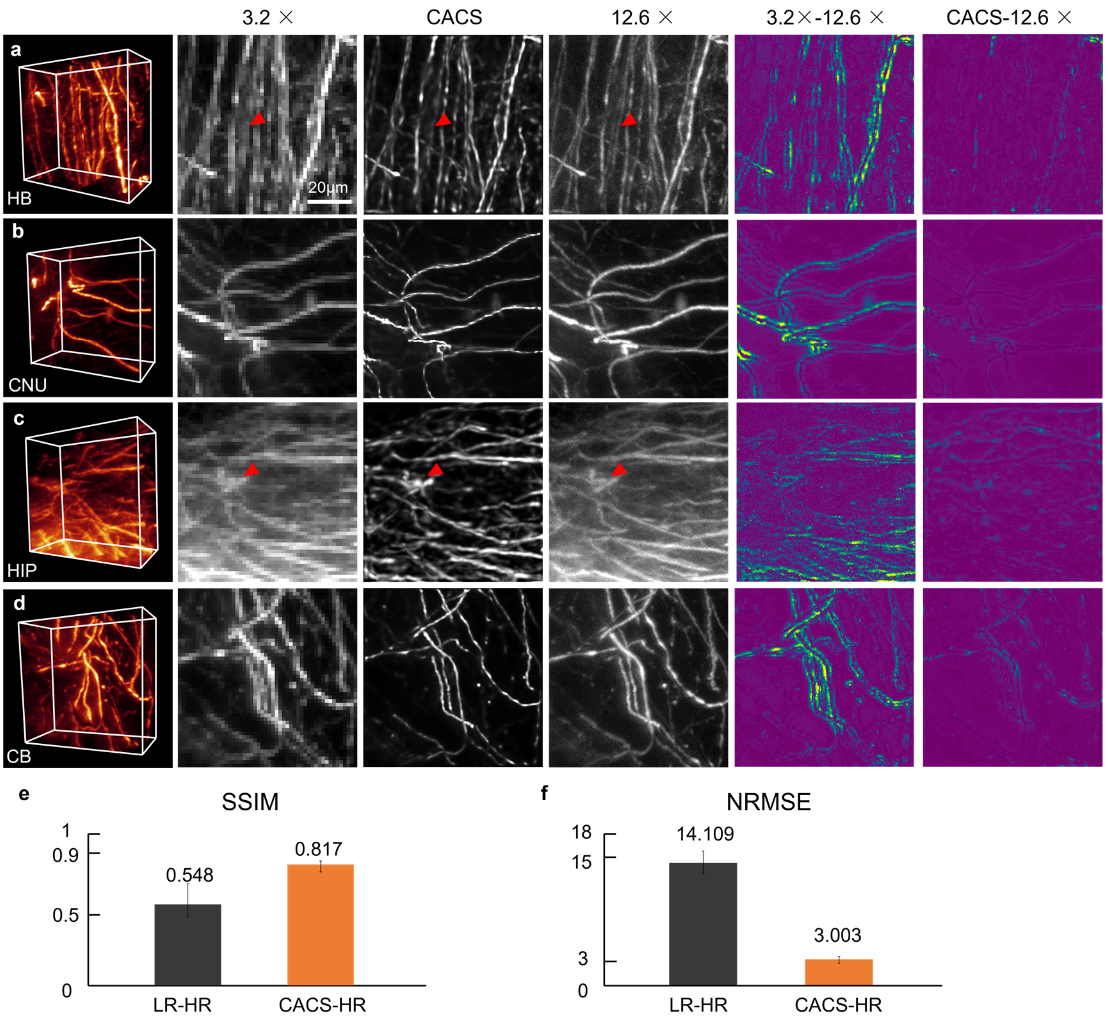


**Supplementary Fig. 6 | CACS computation for line-like neuron signals at different density.** Four ROIs of whole mouse brain (Thy1-GFP-M) containing neuronal fibers with different signal densities are imaged using 3.2× CACS Bessel sheet. 12.6× Bessel sheet results are regarded as ground truth to verify the accuracy of CACS recovery. **a-d**, The comparisons between 3.2×, 3.2× CACS and 12.6× results of the four ROIs in HB, CNU, HIP and CB brain regions, respectively. The red arrows in the ***x-y*** planes show few amount of inaccurately resolved signals by CACS, which are more likely to happen in dense signal regions. The error maps of 3.2× and 3.2× CACS shown at the right two columns also confirmed the high reconstruction accuracy of CACS. **e-f**, The notably lower NRMSE and higher SSIM of CACS results as compared to those of raw Bessel sheet results. Therefore, the perceptual assessments together with quantitative analyses verify the high recovery fidelity of CACS for line-like signals when providing resolution improvement. Scale bar, 20 μm.

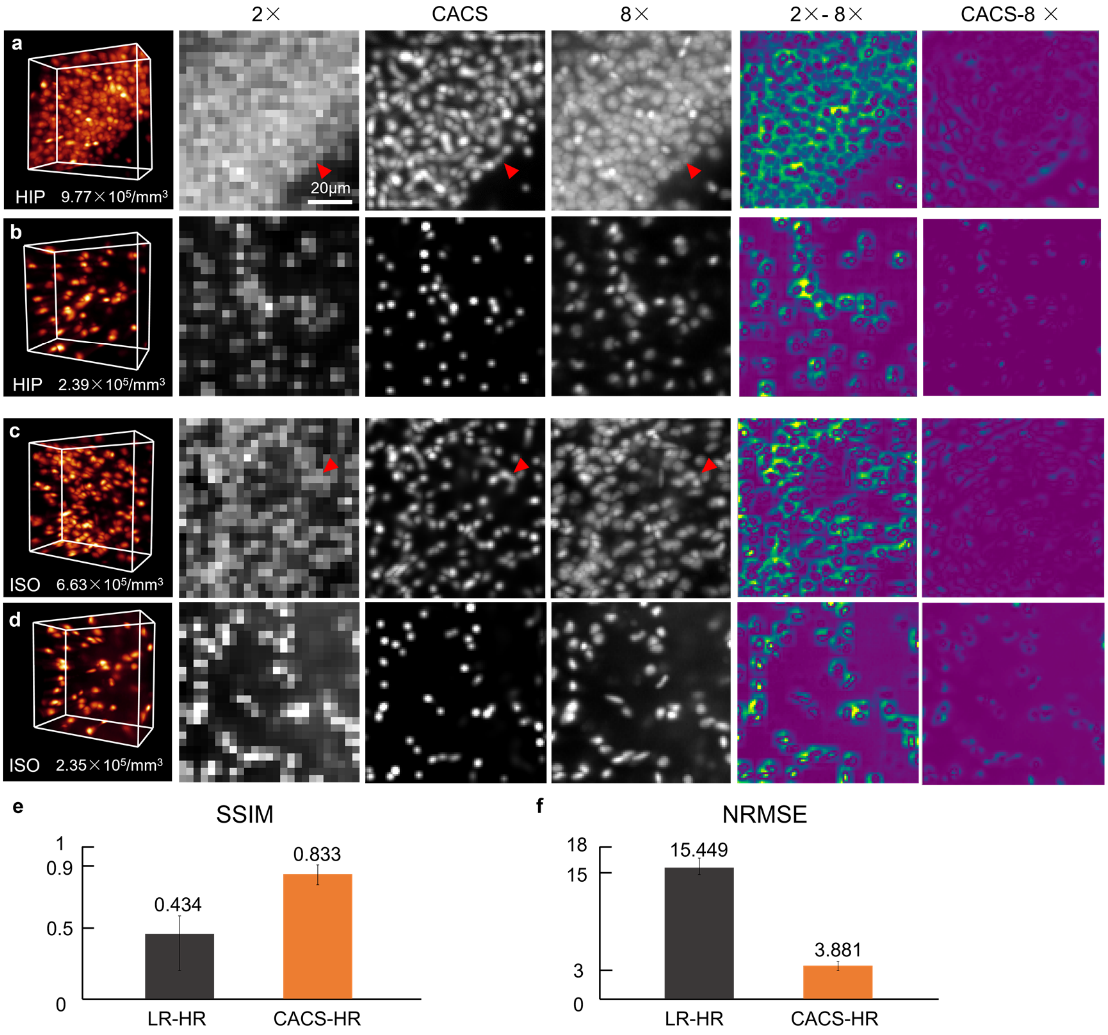


**Supplementary Fig. 7 | CACS computation for point-like cell nuclei at different density. a-d**, The comparisons between 2×, 2× CACS and 8× results of the four ROIs in PI-labelled hippocampus and isocortex, respectively. The ROIs contain cell nuclei with different density. The red arrows in the ***x-y*** planes of **a** and **c** show the inaccurately resolved nuclei signals by CACS, which are also indicated in the error maps at the right columns. **e-f**, The NRMSE and SSIM values of raw 2× images and 2× CACS images. The perceptual assessments together with quantitative analyses all verify the high recovery fidelity of CACS also for point-like signals. Scale bar, 20 μm.

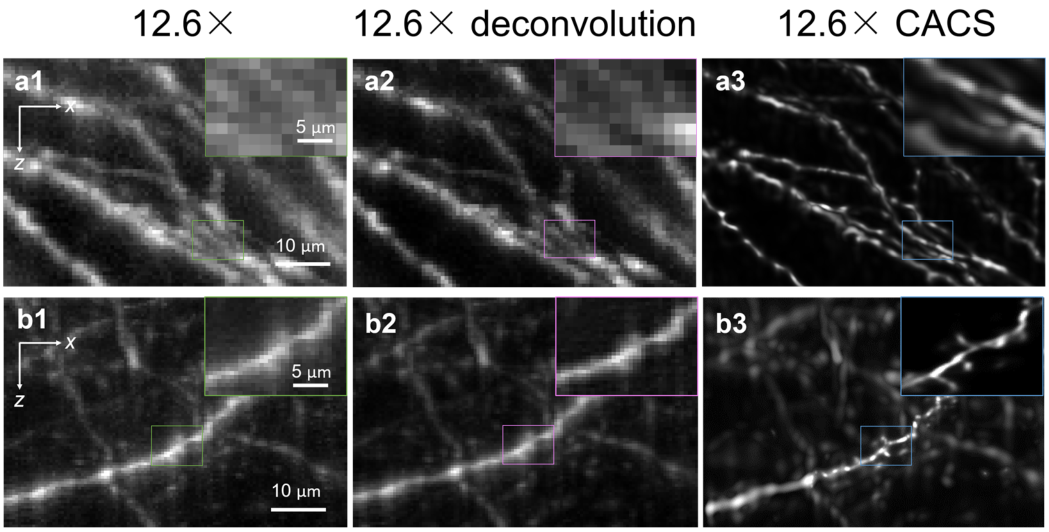


**Supplementary Fig. 8 | CACS computation for resolving finer neuronal sub-structures.** The CACS can be also applied to high-resolution 12.6× Bessel sheet images for super-resolving finer dendrite spines. **a, b,** ***x-z*** planes of two ROIs from isocortex of a Thy1-GFP-M mouse brain. As compared to the raw 12.6× Bessel sheet results **(a1, b1)** and deconvolution results **(a2, b2**), the CACS results reveals the fine structure of the dendrite spines **(a3, b3).** Scale bars, 10 µm (insets, 5 µm).

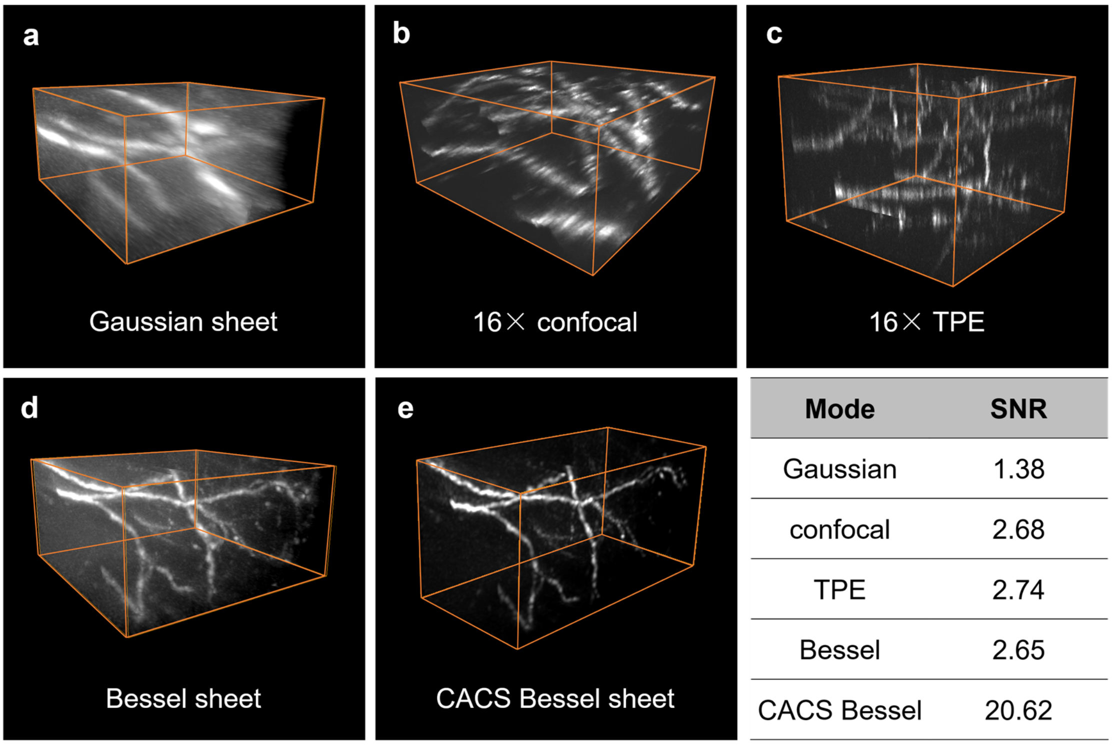


| Mode | SNR |
|---|---|
| Gaussian | 1.38 |
| confocal | 2.68 |
| TPE | 2.74 |
| Bessel | 2.65 |
| CACS Bessel | 20.62 |

**Supplementary Fig. 9 | Signal-to-noise-ratio (SNR) comparison of different modes.** The image SNR by each mode could be obtained via dividing the averaged value of 20% brightest signal voxels by the averaged value of 80% darkest noise voxels. **a-e**, The images of neuronal fibers in cleared brain cortex obtained by Gaussian sheet, 16× confocal, 16× TPE, Bessel sheet, and CACS Bessel sheet, respectively. First, as compared to relatively thick optical sectioning by Gaussian sheet, the thinner and more intensive optical sectioning by Bessel sheet/confocal/TPE yields higher SNR as well as better axial resolution. Then, in addition to resolution improvement, the CS computation also notably increases the SNR of raw Bessel sheet, generating obviously highest-quality image among the five modes. All the images were acquired by each method using parameters listed in **Supplementary Table 1.**

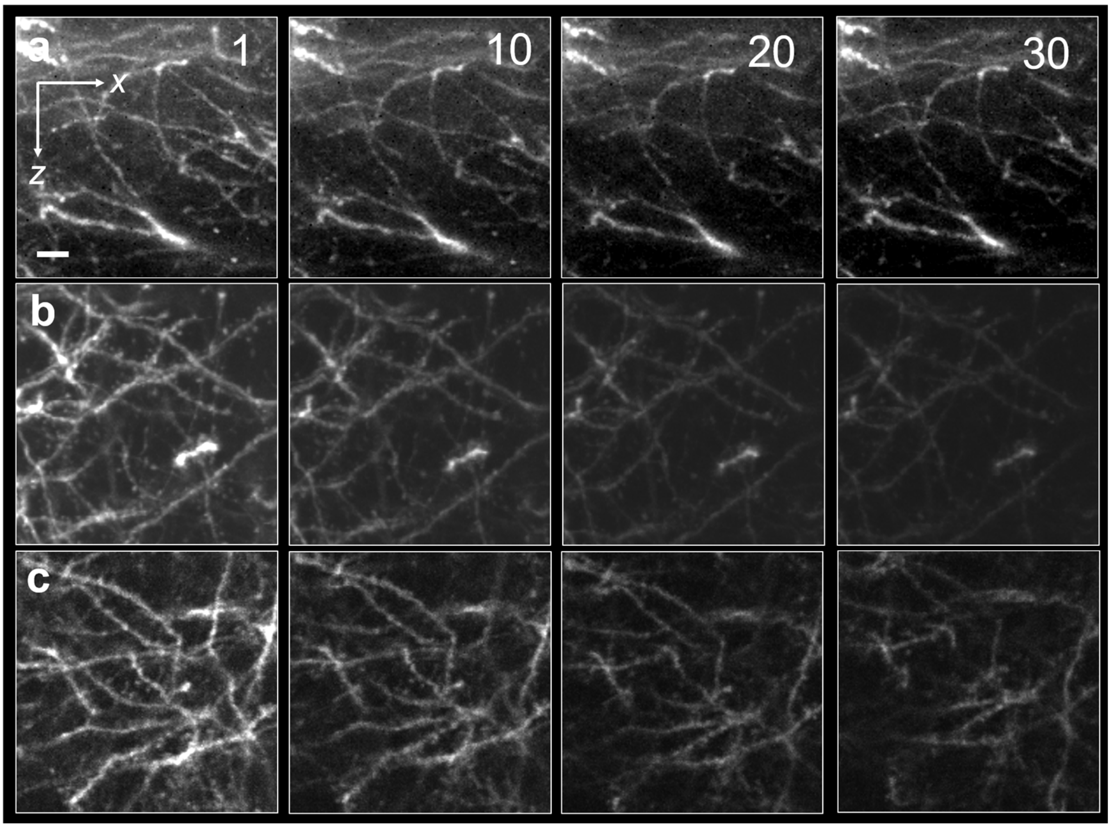


**Supplementary Fig. 10 | Photo-bleaching rate comparisons of different imaging modes.**
**a**-**c**, Maximum-intensity-projections (MIPs) of cortex nerves in the 1$^{st}$,10$^{th}$, 20$^{th}$ and 30$^{th}$ image stacks by Bessel sheet, confocal and TPE, respectively. Scale bar, 5 μm. All the images were acquired by each method using parameters listed in **Supplementary Table 1**.

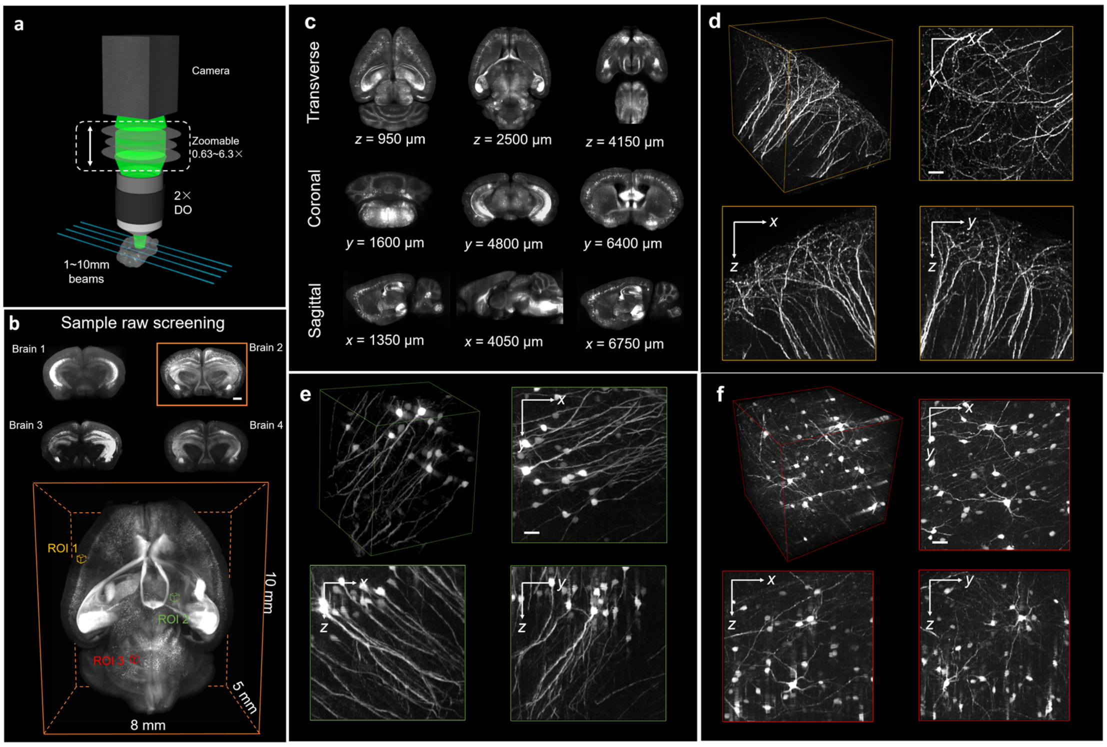


**Supplementary Fig. 11 | Scalable isotropic imaging of neurons in mouse brain. a**, Dual-side, tunable Bessel sheet illumination combined with zoomable detection FOV from 1.26× (~1 cm) to 12.6× (~1 mm). **b**, Reconstructed coronal planes of four mouse brains that are rapidly screened using raw 1.26× Bessel sheet mode. Each brain was imaged under 2 views for generating a multi-view-fused 3D reconstruction that contains 1000 image planes. The bottom row shows volume rendering of a whole brain (No.2) selected, owing to its best signal distribution among 4 candidates. **c**, MIPs in transverse (***x-y***; top), coronal (***x-z***; middle), and sagittal (***y-z***; bottom) planes of the No.2 whole brain, showing the overall signal distributions. Then, higher-resolution imaging of any region of interest was possible using the 12.6× Bessel sheet mode. For example, we imaged three ~1 mm$^3$ regions of interest (ROIs) in the cortex, hippocampus, and cerebellum of brain number 2 at an imaging speed of ~0.01 mm$^3$ s$^{-1}$ and an isotropic resolution of ~1.5 µm (0.5-µm voxel). **d-f**, Three ~1 mm$^3$ ROIs in cortex (yellow), hippocampus (green), and cerebellum (red) regions, selected from the coarse 1.26× reconstruction, and further imaged by 12.6× Bessel sheet mode, to reveal the various neuron types/structures (dense dendrites in **d**, pyramid neurons in **e**, astrocytes in **f**) at subcellular isotropic resolution.

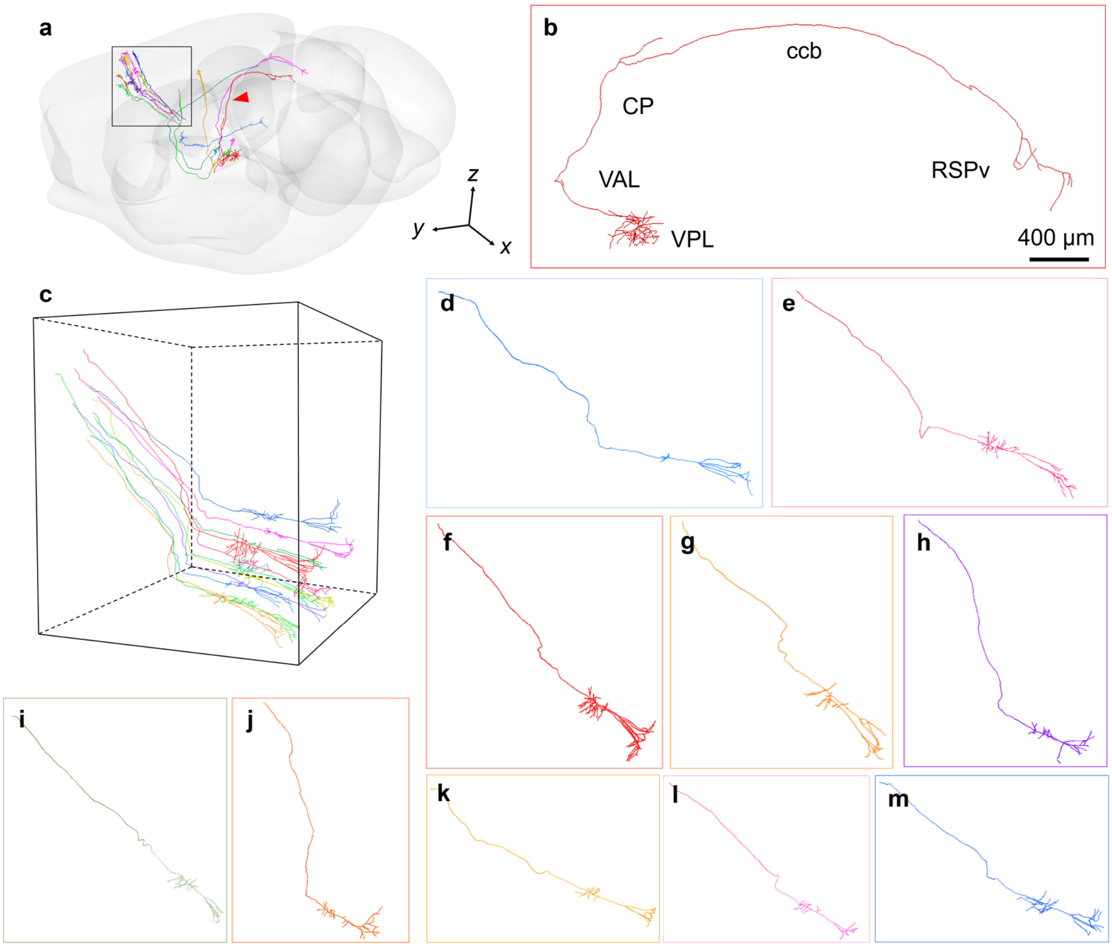


**Supplementary Fig. 12 | Tracing dense long-distance projection neurons in 8-week Thy1-GFP-M mouse brain.** With visualizing whole brain at isotropic subcellular resolution by CACS Bessel sheet, we are able to identify and trace long-distance projection neurons across the whole mouse brain. **a**, Three-dimensional visualization of whole brain with showing the trajectories of long-distance projection neurons. **b**, An annotated projection neuron with pathway across Ventral posterolateral nucleus of the thalamus (VPL), Ventral anterior-lateral complex of the thalamus (VAL), Caudoputamen (CP), corpus callosum, body (ccb), Retrosplenial area, ventral part (RSPv) regions. **c**, Selected cortex area (2 × 2 × 2 $mm^3$ volume) containing a dense bundle of pyramidal tract neurons. 10 projection neurons (**d**-**m)** initiated from this bundle. Thus, the segmentation and tracing of these densely packed neurons intrinsically require high volumetric resolution at large scale.

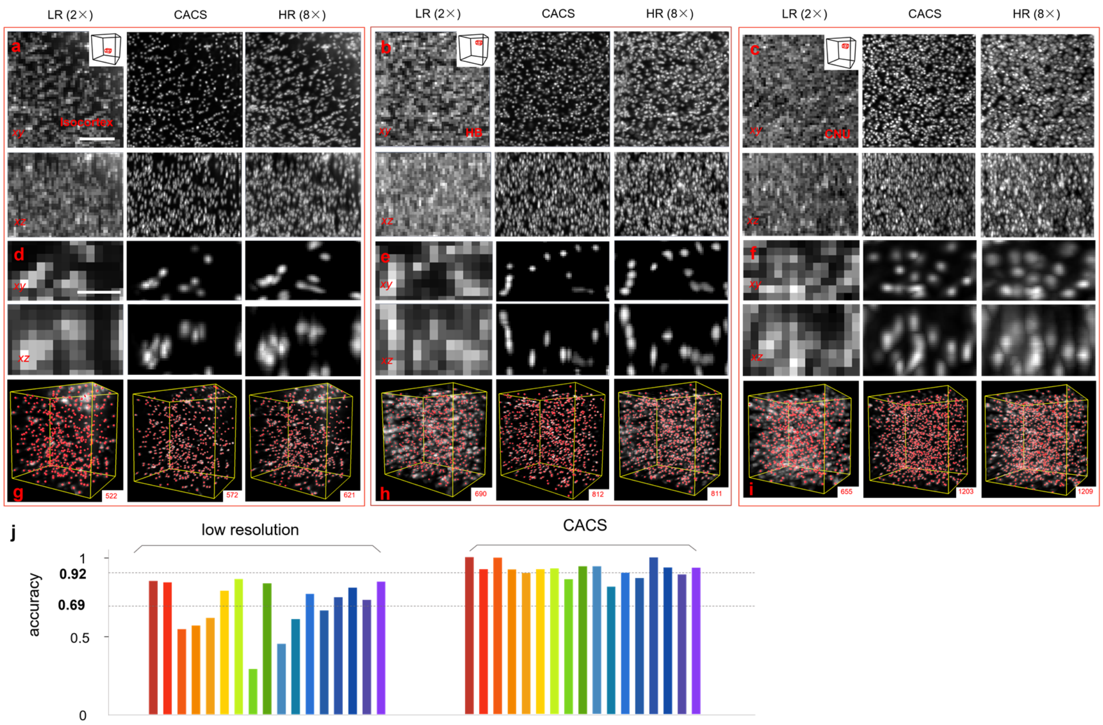


**Supplementary Fig. 13 | Accuracy of compressed sensing in PI-labelled brain.** We verified the high accuracy of CACS-enabled cell counting in seventeen 3D volumes from all the 10 sub-regions of a half brain. **a**-**f**, PI-labelled nuclei images (MIPs) of three selected volumes ($1 \times 1 \times 1$ mm$^3$) from isocortex, HB, and CNU, respectively (**a**-**c**). The signal density in these three sub-regions is different, thus to test the robustness of CACS recovery. Each region was visualized by 2× Bessel sheet, 2× CACS Bessel sheet and 8× Bessel sheet, as shown in left, middle and right column, respectively. The magnified details ($32 \times 60 \times 32$ μm$^3$) from these sub-regions by three modes are compared to highlight the notable resolution improvement of CACS recovery (comparing **d1**-**f1** and **d2**-**f2**), which is known to be highly relevant with subsequent quantitative analyses. At the same time, the recovered signals by 2× CACS were also verified to be sufficiently accurate (comparing **d2**-**f2** and **d3**-**f3**). **g**-**i**, Corresponding nuclei segmentation in 2×, 2× CACS, and 8× images using Imaris, showing different results orientated by the image quality. **j**, The cell counting accuracy, defined as (2× number / 8× number) × 100% or (CACS number / 8× number) × 100%, for all seventeen 3D volumes by 2× Bessel sheet (left) and 2× Bessel-CACS (right). As compared to the standard counting results based on 8× images, the averaged counting accuracy based on 2× results for these volumes of distinct cell density is merely 0.69, while this value is increased to ~0.92, within the error tolerance, based on 2× CACS results. Scale bar, 50 μm.

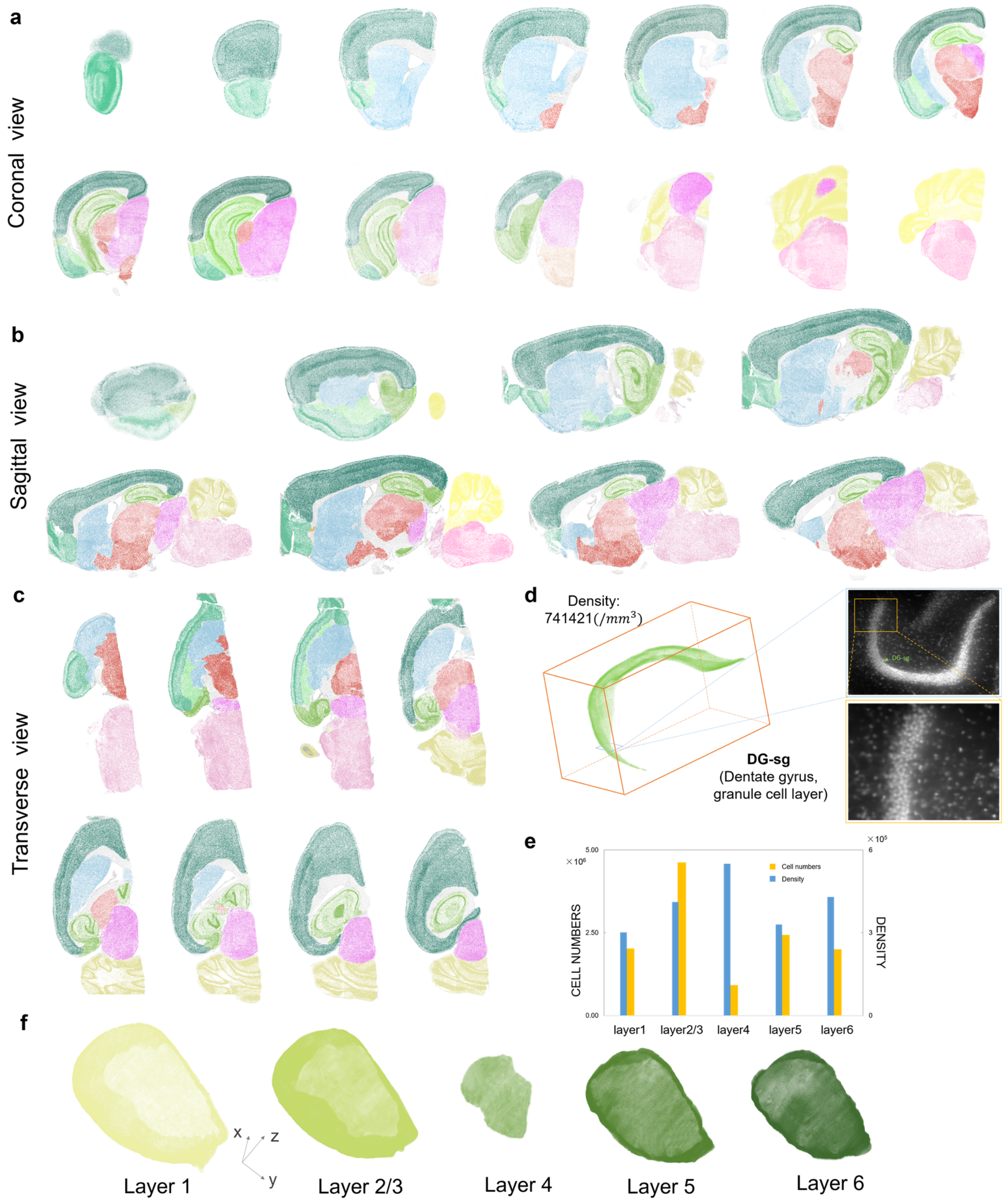


**Supplementary Fig. 14 | Segmentation and cell counting for PI-labelled half brain imaged by 2× CACS Bessel sheet.** With verifying the counting accuracy of CACS Bessel sheet, we applied it to the quantitative analysis of half brain. **a-c**, Series of coronal views, sagittal views and transverse views in different $\boldsymbol{y}$, $\boldsymbol{x}$ and $\boldsymbol{z}$ positions, showing the segmented encephalic regions in a PI-labelled half brain. **d**, Selected 3D reconstruction and image details of granule cell layer at Dentate gyrus. **e**, **f**, Cell numbers and density of different layers in isocortex.

#### Supplementary Note 1 | Imaging speed, photobleaching rate and SNR

The performances compared between the mentioned methods are listed in following table. All the comparison is based on the imaging results of mouse brain.

The speed for each method is calculated in following form:

$$Imaging\ speed\ =\ \frac{Imaging\ volume}{Acquisition\ time}\ (mm^3/second) \qquad (1)$$

We applied a mask to remove all frequency components beyond the Abbe limit in Fourier domain to divide the raw 3D stack to a filtered stack $I_1$ and a noise stack $I_2$. Then the SNR is calculated by

$$SNR = \frac{mean\ (Top_{0.2} I_1)}{RMS\ (Top_{0.2} I_2)} \qquad (2)$$

The bleaching rate curve displays the relationship between imaging times and normalized signal intensity S. A selected 100 µm × 100 µm × 100 µm cortex volume was repetitively imaged 30 times to compare the photobleaching rates. The signal bleaching over the time for each method is shown in **Supplementary Fig. 10.**

The imaging throughput is calculated by

$$Imaging\ throughput = \frac{Number\ of\ voxels}{Acquisition\ time}\ (voxels/second) \qquad (3)$$

**Supplementary Note Table 1 | Comparison of different imaging modes**

<table>
<tr><td colspan="2">Modes</td><td>TPE</td><td>Confocal</td><td>3.2× Gauss</td><td>3.2× Bessel</td><td>3.2× CS</td><td>12.6× Bessel</td></tr>
<tr><td rowspan="4">Imaging parameters</td><td>Magnification/NA</td><td colspan="2">16×/0.8 (Water)</td><td colspan="3">3.2×/0.28 (Air)</td><td>12.6×/0.5 (Air)</td></tr>
<tr><td>Frame rate (fps)</td><td colspan="2">1</td><td colspan="3">40</td><td>20</td></tr>
<tr><td>Pixel size (µm)</td><td colspan="2">0.31 × 0.31</td><td colspan="2">2.03 × 2.03</td><td>(0.508 × 0.508)</td><td>0.516 × 0.516</td></tr>
<tr><td>Z step (µm)</td><td colspan="2">2.5</td><td>4</td><td>2</td><td>(0.5)</td><td>0.5</td></tr>
<tr><td rowspan="3">Acquisition time</td><td>Speed (×10$^6$ µm$^3$/s)</td><td>0.042</td><td>0.042</td><td>2768.89</td><td>1384.37</td><td>1384.37</td><td>11.17</td></tr>
<tr><td>Throughput (×10$^6$ voxels/s)</td><td colspan="2">0.177</td><td colspan="2">157.28</td><td>10737.42</td><td>83.88</td></tr>
<tr><td>Photobleaching</td><td>60%</td><td>70%</td><td>5%</td><td colspan="2">10%</td><td>20%</td></tr>
<tr><td rowspan="2">3D resolution</td><td>Lateral</td><td>0.85</td><td>0.85</td><td colspan="2">4.5</td><td>1.5</td><td>1.5</td></tr>
<tr><td>Axial</td><td>4</td><td>4.2</td><td>15</td><td>4.5</td><td>1.5</td><td>1.5</td></tr>
</table>

**Supplementary Note 2 | Image stitching and dual-view image fusion**

Cleared mouse brain (PEGASOS), or other large organs, still show non-negligible light attenuation and scattering at the deep of tissue. For dual-view 3.2× Bessel imaging of whole mouse brain, we only acquired information at 0-3 mm depth of the brain (totally ~5 mm in depth) for each view, discarding degraded signals from the deepest tissues and thereby reducing the imaging time by ~40%. 6 lateral tiles (4.16 × 4.16 × 3 mm FOV, ~40 gigavoxels) were stitched under each view, to form dual-view whole brain data. Then the complete whole-brain information could be obtained by a bead-based registration of 2 views followed by a weighted image fusion (**Supplementary Table 2**). The registered-and-fused Bessel brain was further processed by CACS to obtain the final digital whole brain with large volume size and high spatial resolution. The implementation details of the whole-brain imaging are listed below (**Supplementary Table 2**). It is noteworthy that 3.2× CACS Bessel sheet can provide subcellular resolution similar with that by 12.6× Bessel sheet while significantly reduce the acquisition time down to ~10 minutes, over 100-folds shorter than the time for 12.6× Bessel sheet.

**Supplementary Note Table 2 | Whole-brain imaging with different magnification**

| Modes | 1.26× Gauss | 1.26× Bessel | 3.2× Bessel | 3.2× CACS Bessel | 12.6× Bessel |
|---|---|---|---|---|---|
| Stitching tiles | 1 | | 2 × 3 | | 8 × 12 |
| Pixel size (μm) | 5.16 | | 2.03125 | 0.5078 | 0.516 |
| Z step (μm) | 4 | 2.5 | 2 | 0.5 | 0.5 |
| Frame rate | 40 | | | | 20 |
| Acquisition speed | 40 s/ whole brain | 60 s/ whole brain | 10 min/ whole brain | | 16.5 h/ whole brain |
| Data storage | 9.76 Gb | 15.625 Gb | 103.9 Gb | 6.49 Tb | 6.07 Tb |

**Supplementary Note 3 | Content aware regularization in CS**

In CACS, a crucial step is to calculate the parameter $\lambda_i$ that indicates image contents in each input image stack $y_i$, using equation: $\lambda_i = \alpha_i \cdot \beta_i$. Here $\alpha_i = \|2A^T y_i\|$ is a parameter calculated by the PSF and signal density in raw image $y_i$, with larger $\alpha_i$ denoting more sparse signals in $y_i$. $\beta_i = kE_i + b$, is another parameter calculated by entropy of raw images, where $E$ term represents the entropy ($E$) of $y_i$ and indicates the degree of signal disorder (**Supplementary Note Fig. 1a, b**). For line-like structures, such as neurons, $k_{line}$ = -0.08 and $b_{line}$ = 0.52; for point-like signals, such as PI-labelled nuclei, $k_{point}$ = -0.35 and $b_{point}$ = 1.82 (**Supplementary Note Fig. 1c**). Thus, the value of $\beta_i$ is inversely proportional to the entropy of the image, with larger $\beta_i$ denoting less-disordered signals in $y_i$. The value of $\lambda_i$ is then determined by the product of these 2 indices, and applied to the regularization term to determine the weight of origin image when algorithm tries to solve the following equations:

$$\|X_{i,n+1}\|_1 = \|AX_{i,n} - Y_i\|_2^2 + \lambda_i \|X_{i,n}\|_1 \quad (4)$$

In traditional CS implementation without content-aware regularization, the recovery of dense signals tends to have over-fitting issue caused by excessive initial constraints, thus showing poor improvement. On the contrary, the recovery of sparse signals require more constraints to prevent too sharp artefacts[1]. Because the density of signals across a large FOV varies dramatically, there are obvious artefacts and signal-loss in the final stitched result (**Supplementary Note Fig. 1e**). Such a content-aware regularization factor $\lambda_i$ in our CACS can properly identify the signal characteristics and carefully balances the results between these two extremes, thereby recovering relatively accurate signals with substantial resolution improvement (**Supplementary Note Fig. 1f, g**). For example, as the calculated $\lambda_i$ being large, also meaning sparse and ordered signals presented in raw image, the algorithm correspondingly has large weighting from the raw image and tends to preserve more existing real information rather than inference of new information.

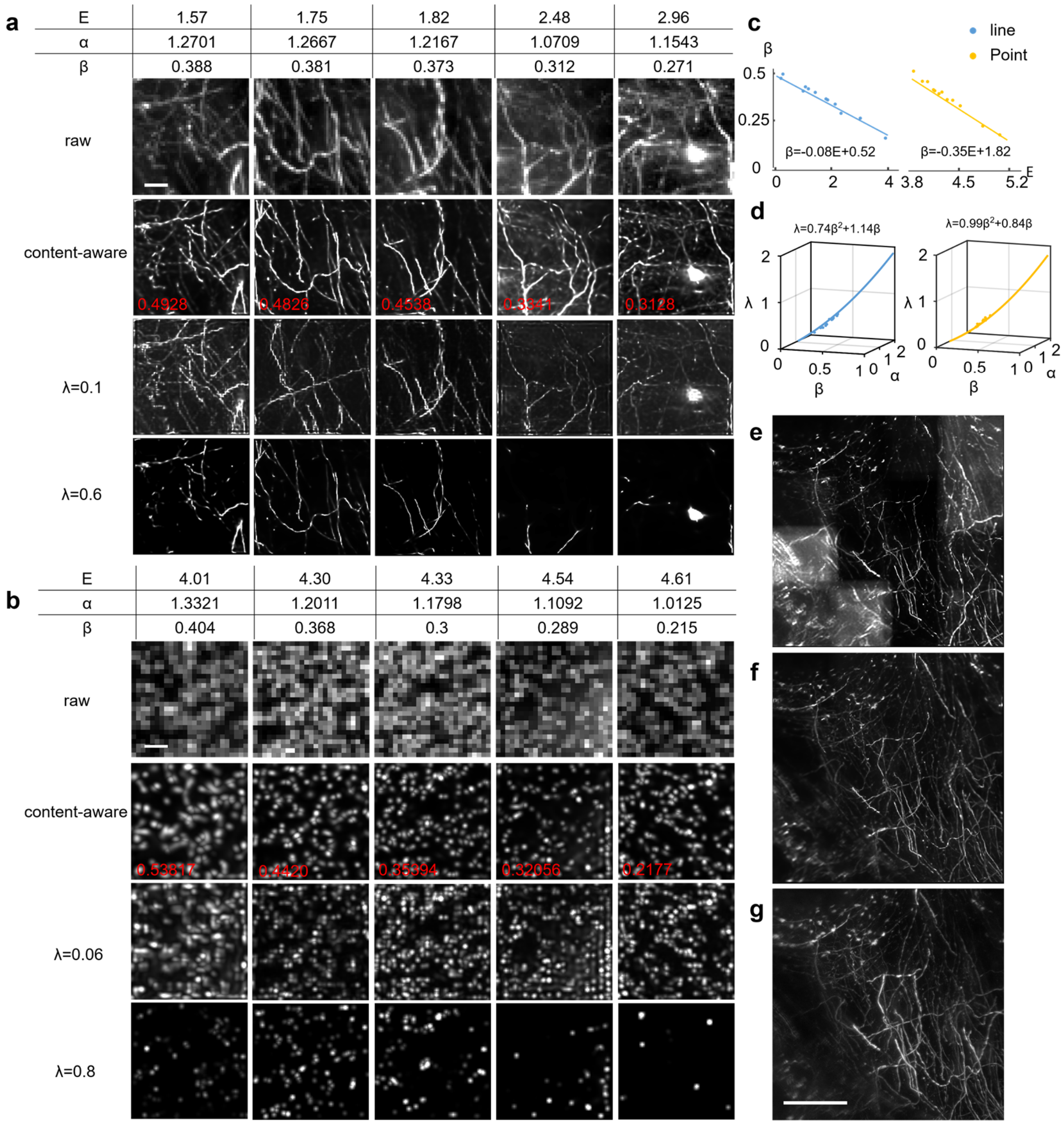


**Supplementary Note Fig. 1 | Content-aware calculation of regularization factor. a**, 5 line-like structures from Thy1-GFP-M mouse brain with applying different sparsity indicator $\boldsymbol{\alpha}$ and entropy indicator $\boldsymbol{\beta}$. Three types of CS enhancement results with $\lambda_1 = \alpha \cdot \beta$, $\lambda_2 = 0.1$ and $\lambda_3 = 0.6$ are compared to show the artefacts for $\lambda_2$, information loss for $\lambda_3$, and obviously higher accuracy by CACS. **b**, 5 point-like structures selected from PI-labelled mouse brain also verify the advantage of CACS. Scale bar, 20 µm. **c,** The plot of parameter $\boldsymbol{\beta}$ versus the entropy of signals ($\boldsymbol{E}$). The fitted curves (solid lines) reveal the inversely proportional relationship between $\boldsymbol{\beta}$ and the entropy, in both point- and line-like signals. **d**, The data plot (scattered points) of regularization fact $\boldsymbol{\lambda}$ versus parameters $\boldsymbol{\alpha}$ and $\boldsymbol{\beta}$. The fitted curves (solid lines) indicate the positive correlation between $\boldsymbol{\alpha}$ and $\boldsymbol{\beta}$, also quasi-quadratic relationship between final $\boldsymbol{\lambda}$ and $\boldsymbol{\beta}$. **e-g**, Comparison of conventional CS computation with constant parameters (**e**), our CACS with adaptive parameters (**f**), and 12.6× ground-truth image (**g**). Scale bar, 100 µm.
